# Response eQTLs, chromatin accessibility, and 3D chromatin structure in chondrocytes provide mechanistic insight into osteoarthritis risk

**DOI:** 10.1101/2024.05.05.592567

**Authors:** Nicole E Kramer, Seyoun Byun, Philip Coryell, Susan D’Costa, Eliza Thulson, HyunAh Kim, Sylvie M Parkus, Marielle L Bond, Emma R Klein, Jacqueline Shine, Susanna Chubinskaya, Michael I Love, Karen L Mohlke, Brian O Diekman, Richard F Loeser, Douglas H Phanstiel

## Abstract

Osteoarthritis (OA) poses a significant healthcare burden with limited treatment options. While genome-wide association studies (GWAS) have identified over 100 OA-associated loci, translating these findings into therapeutic targets remains challenging. Integrating expression quantitative trait loci (eQTL), 3D chromatin structure, and other genomic approaches with OA GWAS data offers a promising approach to elucidate disease mechanisms; however, comprehensive eQTL maps in OA-relevant tissues and conditions remain scarce. We mapped gene expression, chromatin accessibility, and 3D chromatin structure in primary human articular chondrocytes in both resting and OA-mimicking conditions. We identified thousands of differentially expressed genes, including those associated with differences in sex and age. RNA-seq in chondrocytes from 101 donors across two conditions uncovered 3782 unique eGenes, including 420 that exhibited strong and significant condition-specific effects. Colocalization with OA GWAS signals revealed 13 putative OA risk genes, 10 of which have not been previously identified. Chromatin accessibility and 3D chromatin structure provided insights into the mechanisms and conditional specificity of these variants. Our findings shed light on OA pathogenesis and highlight potential targets for therapeutic development.

**Highlights:**

∘ Comprehensive analysis of sex- and age-related global gene expression in human chondrocytes revealed differences that correlate with osteoarthritis
∘ First response eQTLs in chondrocytes treated with an OA-related stimulus
∘ Deeply sequenced Hi-C in resting and activated chondrocytes helps connect OA risk variants to their putative causal genes
∘ Colocalization analysis reveals 13 (including 10 novel) putative OA risk genes

## Introduction

Osteoarthritis (OA) affects over 500 million individuals globally and is a leading cause of disability in the US^1^; however, treatment options have been elusive in large part because the mechanisms driving OA remain poorly understood. Genome-wide association studies (GWAS) have identified over one hundred OA-associated loci^2,3^. Translating these loci into new knowledge and actionable therapeutic targets requires identification of the genes affected at each GWAS locus, which has proven challenging for multiple reasons. Linkage disequilibrium between nearby variants makes it difficult to identify the causal variant(s) at each locus. Further, most disease-risk variants alter non-coding regulatory sequences, which can affect gene expression over distances exceeding 1 million base pairs, often via 3D chromatin structures that bring those regulatory loci into close physical proximity with their target genes. Finally, these regulatory mechanisms are dynamic across cell types and biological conditions^4,5^; therefore, understanding the functional impact of risk variants in the relevant cellular context is essential. Despite these challenges, coupling genomic and genetic technologies to the appropriate disease models can overcome these hurdles and reveal disease-risk genes for further research and therapeutic development.

Expression quantitative trait loci (eQTL) mapping is a powerful technique for identifying the genes mediating disease risk at each GWAS locus as it directly connects genetic variants to differences in gene expression across a cohort of donors^6,7^. Once mapped, colocalization of these eQTLs with GWAS data can reveal the gene expression changes that likely influence disease risk^8^. The integration of colocalized eQTLs with other genomic datasets including Hi-C and ATAC-seq can provide further support and mechanistic insight into the disease-causing mechanisms at these loci^9^. The power of QTL mapping has fueled large consortiums to generate QTLs for a broad array of tissues^10^ and led to breakthroughs in our understanding of several diseases, including Alzheimer’s disease^11^ and various immune-related diseases^12–14^. Notably underrepresented from these studies are maps of eQTLs in human chondrocytes, which are the only cell type in cartilage, the most OA-relevant tissue in the body^15^. To the best of our knowledge, only one eQTL study has been performed in human chondrocytes which used tissue from donors with advanced OA undergoing joint replacement. This study successfully identified 4 colocalized eQTL/GWAS signals and 1 pQTL/GWAS signal pointing to 5 putative OA risk genes^16^. This was a breakthrough study for OA but leaves a large portion of the 100 OA-associated loci unexplained.

It has become increasingly clear that to understand human disease, it is critical to map QTLs in the correct cell type and biological context^17^. OA risk variants are likely to impact chondrocyte function since cartilage degradation and loss is a central feature of OA and OA risk variants are enriched in chondrocyte regulatory elements. We have previously shown that OA GWAS variants are enriched in chondrocyte regulatory regions suggesting that many OA risk variants likely impact chondrocyte function^18^. To understand the mechanisms that contribute to OA, we have previously established an *ex vivo* model of the OA chondrocyte phenotype using primary human articular chondrocytes. Chondrocytes isolated from normal cartilage obtained from cadaveric human tissue donors^19^ are grown in culture and treated with physiological levels of a fibronectin fragment (FN-f), a cartilage matrix breakdown product, found in OA cartilage and synovial fluid, that triggers changes in cell signaling and gene expression that are characteristic of changes observed in chondrocytes isolated from OA tissue^20–22^. This system is ideal for dissecting OA GWAS signals because (1) it uses primary human cells that are representative of the disease process, (2) studying the response to a controlled stimulus found in OA decreases variability found in OA tissue and allows for the study of earlier stages of the disease, and (3) the *ex vivo* nature of the system allows for functional follow up experiments.

We mapped gene expression, chromatin accessibility, and 3D chromatin structure in primary human chondrocytes treated for 18 hours with either PBS (control) or a purified recombinant fibronectin fragment (FN7-10) that binds to and activates the α5β1 integrin^23^. We identified sex-, age-, and treatment-related changes in chondrocyte gene expression, and intersected those with changes observed in OA tissue providing new insights into how these risk factors influence the OA phenotype. We performed eQTL analysis in both PBS and FN-f-treated conditions revealing thousands of eSNP-eGene pairs, hundreds of which were specific to only one of the two conditions. Colocalization of these signals with OA GWAS data revealed 13 putative OA risk genes, 10 of which had not been implicated by prior eQTL studies. We mapped chromatin accessibility to prioritize putative causal variants within these loci and 3D chromatin structure to offer further support and mechanistic insight for several of these colocalized signals. One gene implicated by these analyses was *PAPPA*, which was upregulated in OA tissue, upregulated in response to FN-f, upregulated with age, and is characterized by a chromatin loop that connects its promoter to GWAS variants over 400 Kb away. This study is a critical step forward for OA as it has identified 10 novel putative OA risk genes for further research and therapeutic development.

## Results

### FN-f induces OA-like transcriptional changes in primary human chondrocytes

To determine how FN-f impacts transcription in human chondrocytes, we performed RNA-seq on chondrocytes from 101 donors. We isolated postmortem human articular chondrocytes from deceased human tissue donors through enzymatic digestion of cartilage tissue, and treated cells for 18 hours with either PBS or with 1μM recombinant FN-f within one week of isolation^23^. We performed high-quality RNA-seq to an average depth of 101.8 million stranded paired-end reads per library (**Fig S1A,B**). Sample gene expressions clustered primarily by treatment after principal component analysis (PCA) (**Fig S1C**). We performed technical replicates on 3 donors and demonstrated a higher correlation between replicates than between different donors (**Fig S1D**). Differential expression comparing FN-f to PBS-treated samples revealed 1850 and 2076 up and downregulated genes, respectively (DESeq2^24^, adjusted p < 0.05, absolute log2 fold change > 1), including more stringently defined sets of 857 and 578 up and downregulated genes, respectively, that exhibited the largest and most significant changes (**Fig 1A, Table S1;** DESeq2^24^, adjusted p < 0.01, absolute log2 fold change > 2). Upregulated genes were enriched for GO terms and KEGG pathways consistent with an OA phenotype including “collagen catabolic process”, “acute inflammatory response”, and “NF-kappa B signaling pathway” (**Fig 1B, Table S2**). Upregulated genes included many that have been previously implicated in OA including *IL1B, MMP13*, and *NFKB*. The promoters of upregulated genes were also enriched for transcription factor (TF) binding motifs for proteins implicated in OA including NFKB and members of the AP-1 complex (**Fig 1C left;** HOMER^25^, p < 0.001). The members of those transcription factor complexes showed concordant changes with the motif enrichment analyses (**Fig 1C right**), which further supports the role these TFs play in the transcriptional response to FN-f. Genes that had previously been shown to be up and downregulated in OA tissue showed the same directional changes in response to FN-f suggesting that our *ex vivo* system is a reasonable model of the OA phenotype (**Fig 1D**; Wilcox test, p < 0.01).

**Fig 1:**
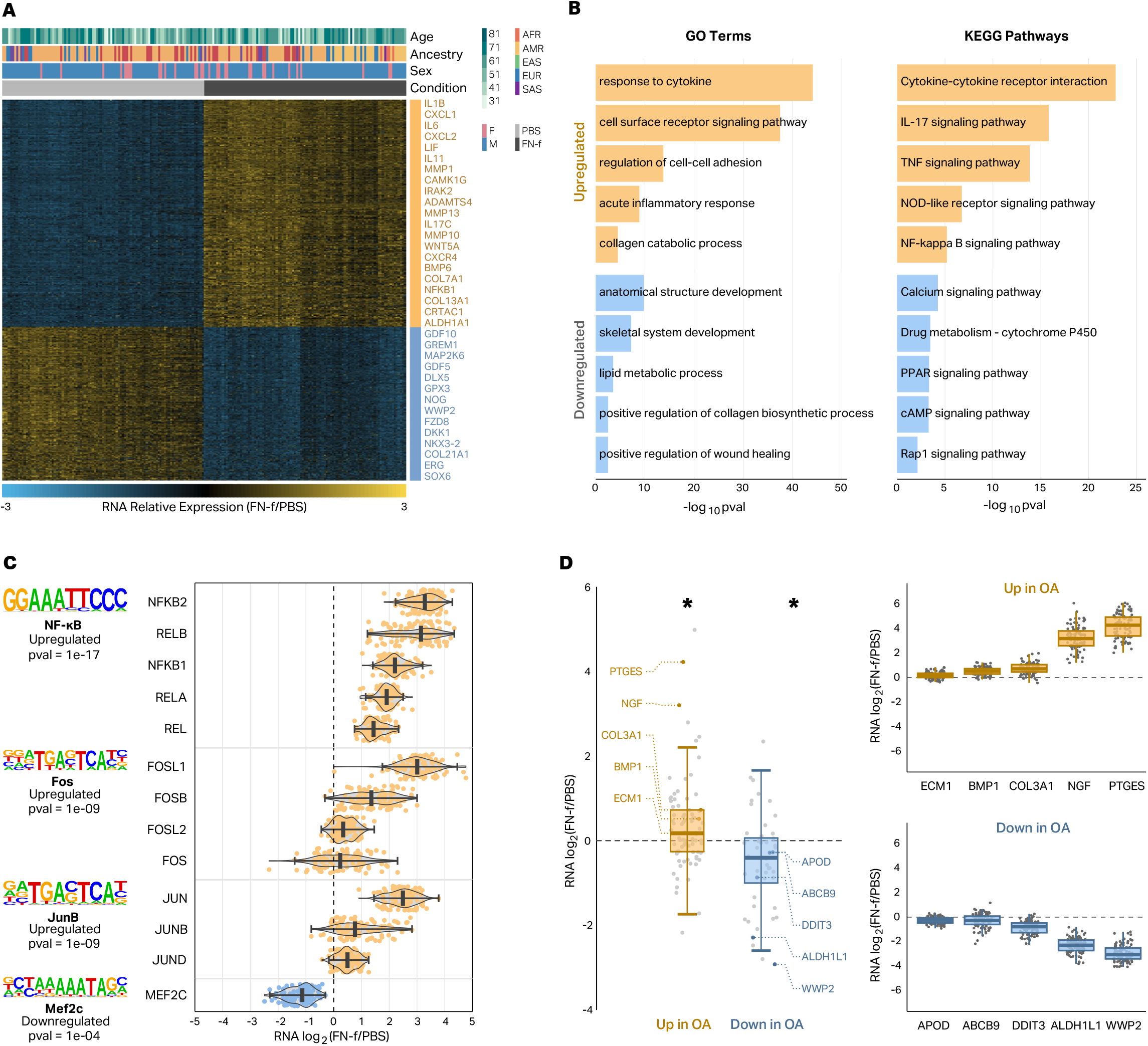
FN-f induces OA-like transcriptional changes in primary human chondrocytes. (**A**) Heatmap depicting the stringent set of 1435 genes differentially expressed between chondrocytes treated with either PBS or FN-f (DESeq2, adjusted p < 0.01, absolute log2 fold change > 2). A subset of the differential genes is listed to the right of the heatmap with upregulated genes colored in yellow and downregulated genes colored in blue. (**B**) Barplots depicting select GO terms and KEGG pathways enriched in sets of genes that are up (yellow) or downregulated (blue) in response to FN-f. (**C**) Names and transcription factor motif logos (left) of transcription factors whose motifs are enriched at the promoters of genes that were up or downregulated in response to FN-f. Violin plots (right) depicting the RNA log2 fold change of members of each transcription factor complex exhibit good agreement with the motif analyses. (**D**) Boxplots (left) show that genes that are up or downregulated in OA tissue show the same trends in chondrocytes treated with FN-f. * indicates Wilcox test p-value < 0.01. Donor-level RNA log2 fold change boxplots highlight examples of genes upregulated (top right) or downregulated (bottom right) in OA tissue.

### Genes with sex- and age-dependent expression patterns include OA-related genes

OA is characterized by sex-related differences in disease risk and severity^26^. These differences could be driven in part by sexual dimorphism in chondrocyte gene expression, either at baseline levels or in response to cartilage matrix damage. Previous studies have investigated sex differences in chondrogenic progenitor cells^27^ and human chondrocytes ^28,29^ from OA tissue, but a comprehensive analyses of sex-related differences in chondrocytes from non OA tissue or in response to cartilage matrix damage have not be conducted. To determine how sex impacted chondrocyte gene expression and if any of these differences corresponded to changes seen in OA tissue, we identified differential expression between sexes in PBS and FN-f-treated samples, while controlling for differences in age and genetic ancestry. We identified 108 genes that differed significantly between sexes (**Fig 2A, Table S3;** DESeq2; adjusted p < 0.01). Most, but not all of these genes were located on chromosomes X and Y (**Fig S2A**). The majority (70%) of sex-related differences in expression were only identified in one condition (**Fig S2B**) and the genes that exhibited sex-related expression differences in both conditions all showed the same direction of effect. Comparison to sex-related expression differences previously identified across 44 tissues from the GTEx consortium^30^ revealed that 29.6% of the sex differences that we observed were unique to chondrocytes (**Fig 2B**, left, **Table S4**). Interestingly, these included the genes that showed the largest fold changes between sexes (**Fig 2B**, right). 35 sex-related genes observed in chondrocytes were also previously shown to be differentially expressed in OA tissue which could provide clues into the sex-related differences in the prevalence and phenotypic presentation of OA (**Table S3**). Examples of sex-related genes either upregulated or downregulated in OA tissue (*SERPINE2* and *RARRES2*) are shown in **Fig 2C**. *SERPINE2* has been shown to inhibit MMP13 in IL1α-treated human chondrocytes^31^. In response to FN-f its expression is higher in male vs female donors, suggesting a stronger protective role in males consistent with the higher prevalence of OA in females^29,32^.

**Fig 2:**
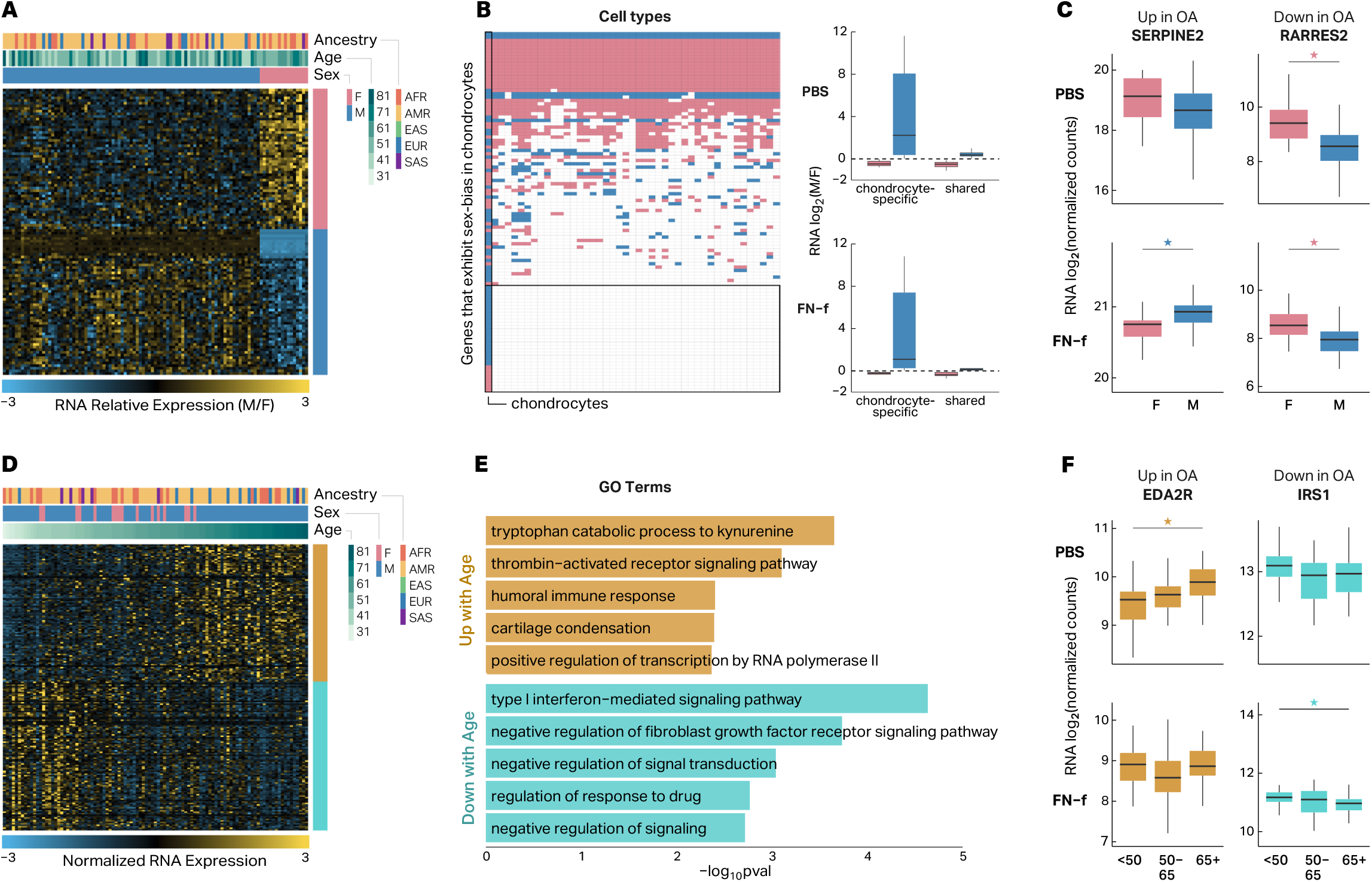
Genes with sex- and age-dependent expression patterns include OA-related genes. (**A**) Heatmap depicting the 108 genes that exhibit sex-biased expression in human chondrocytes. (**B**) Heatmap depicting genes that exhibit sex-biased expression in human chondrocytes and whether or not they exhibit sex-biased expression across 44 other cell lines and tissues analyzed by the GTEx consortium (left). Boxplots depicting the fold-changes (M vs F) of sex-biased genes that are specific to chondrocytes or shared with at least one other tissue (right). (**C**) Boxplots depicting RNA log2 fold change of two genes (*SERPINE2* and *RARRES2*) that exhibit sex-biased expression in human chondrocytes and are differentially expressed in OA tissue. Stars indicate significance (adjusted p-value < 0.01) and are colored by male-biased (blue) or female-biased (pink) expression. (**D**) Heatmap depicting the 196 genes that exhibit age-related expression changes in human chondrocytes. (**E**) GO and KEGG terms that are enriched in genes that show increased (gold) or decreased expression (turquoise) with age in human chondrocytes. (**F**) Boxplots depicting RNA log2 normalized counts of two genes (*EDA2R* and *IRS1*) that exhibit age-related expression changes in human chondrocytes and are differentially expressed in OA tissue. Stars indicate significance (adjusted p-value < 0.05) and are colored by genes that are up with age (gold) and down with age (turquoise).

Age is one of the biggest risk factors for OA, and age-related gene expression in various tissues profiled by the GTEx project has revealed enrichments for genes related to a number of human diseases^33^; however, to the best of our knowledge, there has not been a high-powered analysis of age-dependent RNA-seq-derived gene expression in human chondrocytes. We identified 196 genes that exhibited age-related changes in chondrocyte gene expression (**Fig 2D, Table S5;** DESeq2; adjusted p < 0.05). These genes were enriched for several GO terms and KEGG pathways that are relevant to OA including “cartilage condensation” and “type 1 interferon-mediated signaling pathway” (**Fig 2E, Table S6;** HOMER, p < 0.01). The majority (80%) of age-related changes in gene expression were detected in only one condition (**Fig S2C**), and comparison to age-related gene expression in a subset of GTEx tissues^33^ showed that 99 out of 196 (50.5%) age-related genes were only identified in chondrocytes (**Fig S2D**). 80 of these genes were also previously shown to be differentially expressed in OA vs non-OA tissue, including *EDA2R* and *IRS1* (**Fig 2F**). EDA2R is a member of the TNF receptor superfamily, and the pro-inflammatory TNF pathway has been previously implicated in OA^34^. An increased expression of *EDA2R* in older donors is consistent with “inflammaging” that may contribute to OA pathogenesis^35^. IRS-1 is a mediator of IGF signaling, which has been shown to be reduced in articular chondrocytes in an age-related manner contributing to reduced anabolic activity in cartilage.^36^

### Genetic differences impact gene expression in resting and activated chondrocytes

To determine the impact of genetic differences on chondrocyte gene expression, we performed expression QTL analysis on both PBS- and FN-f-treated samples and tested the association of each gene’s expression with genetic variants within ± 1Mb from the transcription start site (TSS). After hierarchical multiple testing correction with the Storey-Tibshirani q-value^37^ (qval < 0.05), we identified 3782 unique eGenes (**Fig S3**). We then used a conditional analysis (see methods) to identify genes with multiple independent signals and identified 2988 conditionally independent eQTL signals corresponding to 2707 unique eGenes in PBS-treated chondrocytes and 3065 distinct eQTL signals corresponding to 2746 unique eGenes in FN-f-treated chondrocytes (**Table S7**). 267 PBS eGenes and 305 FN-f eGenes had two or more independent signals, including the matrix metalloproteinase *MMP16* (**Fig S4**). Our results captured the majority (64.6%) of the eGenes identified by Steinberg et al.^16^ and increased the total number of eGenes by more than two-fold (3782 vs 1569; **Fig S5A, Table S8**). The effect sizes of the eQTLs for shared eGenes between our study and the lead eQTLs from Steinberg et al. exhibited a strong correlation (mean R^2^ = 0.84; **Fig S5B**). The majority (55.8%) of identified lead eGene-eSNP pairs were only identified in one condition, highlighting the value of mapping eQTLs in specific biological conditions (**Fig 3A, Fig S3C**). By explicitly testing for the interaction between condition and genotype, we identified 696 lead eQTLs with a stronger genetic effect in PBS-treated cells and 856 lead eQTLs that exhibited a stronger genetic effect in FN-f treated cells (i.e. response eQTLs; **Fig 3A**). We further filtered these eQTLs for those that were only identified in one condition, had at least 5 donors with each variant genotype, and had a beta difference of at least 0.2 between conditions, thus producing a refined list of high-confidence PBS-specific eQTLs and FN-f response eQTLs (**Fig 3B-D, Table S9**). Several of these response eQTLs marked genes with known roles in OA including *DIO2* (**Fig 3B**), whose increased expression has been shown to disturb cartilage matrix homeostasis^38^, and *SMAD3*, which, along with the TGF-β signaling pathway, is required for repressing chondrocyte hypertrophic differentiation^39,40^. Several KEGG pathways that are enriched in our set of condition-specific eGenes (**Table S10**) are relevant to OA including apelin signaling and FoxO signaling. Apelin is an adipokine that has been shown to activate catabolic signaling and promote OA progression in preclinical models of OA^41,42^. The FoxO family of transcription factors, including FoxO1, 3, and 4, promote cartilage homeostasis while a decline in FoxO signaling seen in aging and OA is thought to promote cartilage damage^43^.

**Fig 3:**
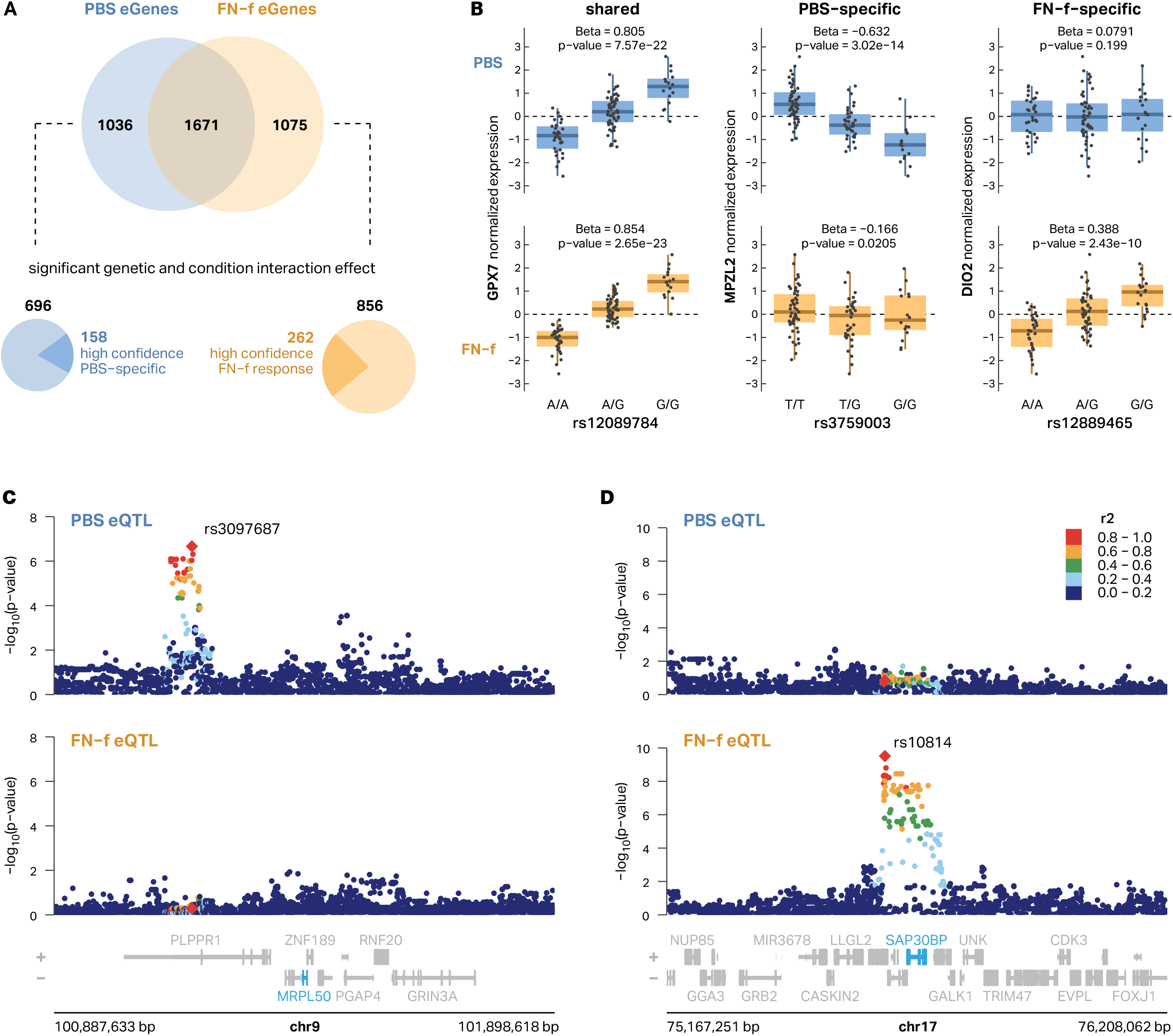
Genetic differences impact gene expression in resting and activated chondrocytes. (**A**) A Venn diagram depicts the overlap between eGenes identified in PBS vs FN-f-treated conditions (top). Explicitly testing for the interaction between condition and genotype we identified 696 lead eQTLs with a stronger effect in PBS-treated cells and 856 lead eQTLs that exhibited a stronger effect in FN-f -treated cells (response eQTLs) which were further filtered to reveal 158 and 262 high-confidence PBS- and FN-f-specific lead eQTLs respectively (bottom). (**B**) Boxplots of genotypes vs normalized expression depicting examples of PBS-specific (*MPZL2*), FN-f-specific (*DIO2*), and shared (*GPX7*) eQTLs. Locus zoom plots show examples of (**C**) PBS-specific (*MRPL50*) and (**D**) FN-f-specific (*SAP30BP*) eQTL signals. Signals are colored by LD to the signal lead variant, denoted as red diamonds and labeled with their rsID.

### Chromatin accessibility supports response eQTLs and refines lists of putative causal variants

To gain insight into the possible mechanisms via which eQTLs exert their effect, we mapped chromatin accessibility using ATAC-seq in chondrocytes from 3 individuals treated with either PBS or FN-f. We identified 217,039 chromatin accessibility peaks, 27,799 of which differed between conditions (DESeq2, adjusted p < 0.01, absolute log2 fold-change > 1; **Table S11**). Of 320,986 distinct eSNPs from either condition, 6.41% of them (20,579) overlapped a chromatin-accessible region. 270 of 379 (71.2%) chromatin-accessible regions that overlapped FN-f-specific lead variants and LD proxies (r^2^ > 0.8) exhibited increased accessibility in FN-f -treated cells (**Fig S6A**) and were enriched for the binding motifs of transcription factors with known roles in chondrocyte matrix damage response including AP-1 (**Fig S6B**). These results further support the validity of our response eQTLs, provide a refined list of variants that might be driving the eQTLs, and point to possible mechanisms through which these variants may act.

### 3D chromatin structure supports distal eSNP-eGene connections

Many of the eSNP-eGene connections we identified suggested long-range regulatory contacts as 24.5% of the lead eSNPs were more than 100 Kb from the nearest promoter of their corresponding eGene (**Fig S7A**) and 44.5% of eSNP-eGene connections ‘skipped’ at least one closer gene (**Fig S7B**). To map potential regulatory connections and determine if 3D chromatin architecture could explain these distal eSNP-eGene pairs, we performed in situ Hi-C in primary human chondrocytes from four donors treated with either PBS or FN-f. We identified 9,099 loops, including 53 that exhibited a significant change in contact frequency between conditions (DESeq2, adjusted p < 0.1; **Table S12**), all of which were increased in contact frequency in response to FN-f. Genes at the anchors of these gained loops included many key players in chondrocyte response to matrix damage and OA including *JUN, IL6*, and *MMP13* (**Fig 4A-C**). Genes at the anchors of gained loops also exhibited significant increases in expression in response to FN-f (median fold-change = 3.7, **Fig 4D**) and were enriched for OA-relevant GO terms including ‘regulation of inflammatory response’, ‘reactive oxygen species metabolic process’, ‘extracellular matrix disassembly’, and ‘regulation of catabolic process’ (**Fig 4E**). Lead eSNPs exhibited stronger contact frequency with their associated eGenes than distance-matched genes (**Fig 4F**) providing a possible mechanism for the distal regulation. Condition-specific eSNP-eGene pairs were associated with stronger contact frequency in the condition associated with the eQTL (**Fig 4G**), suggesting that some of the condition-specific effects could be explained by changes to 3D chromatin structure. **Figure 4H** shows an example of one distal eSNP-eGene pair that is supported by a chromatin loop. SNPs that are in moderate to high LD (r^2^> 0.6) with lead eSNP rs10453229 are linked to eGene *LPAR1* via a 400 Kb chromatin loop. *LPAR1* codes for the lysophosphatidic acid (LPA) receptor. Previous work has shown that LPAR1 signaling is required for development of collagen-induced arthritis in mouse models^44^ and LPA was associated with neuropathic pain in a rat OA model^45^. These data provide a mechanistic explanation for distal eSNP-eGene pairs and further support condition-specific eQTLs.

**Fig 4:**
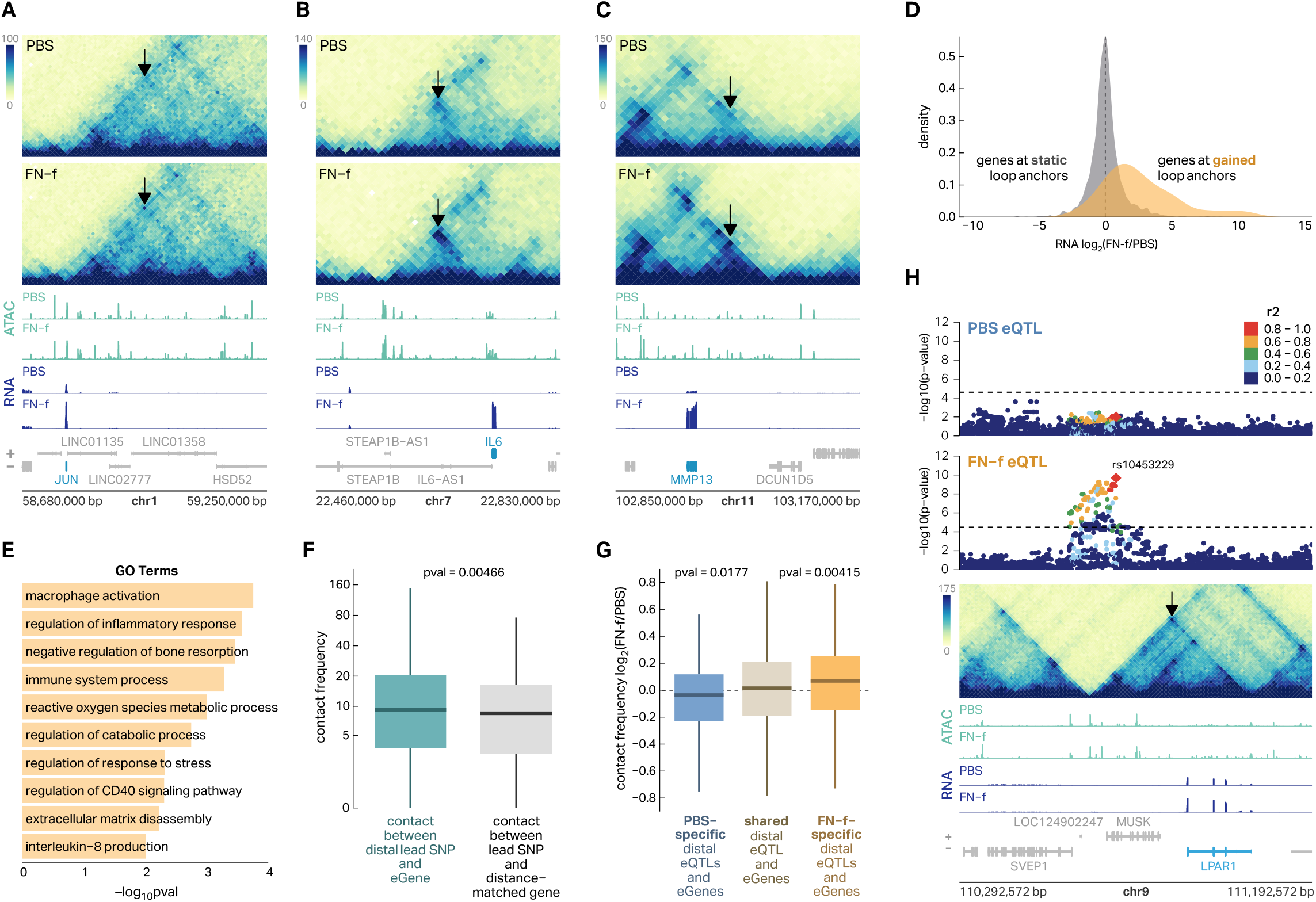
3D chromatin structure supports distal eSNP-eGene connections. (**A–C**) Hi-C, ATAC-seq, and RNA-seq reveal gained enhancer-promoter looping and increased gene expression with FN-f at three OA-associated genes—*JUN, IL6*, and *MMP13*. Arrows point to FN-f-gained loops. (**D**) A density plot of log2 RNA fold-change shows that genes whose promoter overlaps a loop that is gained in response to FN-f also exhibit increases in gene expression. (**E**) A barplot depicting GO terms that are enriched in the set of genes whose promoters overlap gained loop anchors. (**F**) A boxplot shows that lead eSNP-eGene pairs exhibit higher contact frequency than distance-matched pairs of lead eSNPs and genes. P-value was calculated with a Wilcox test. (**G**) A boxplot shows that condition-specific eQTLs were associated with directionally concurrent changes in Hi-C contact frequency suggesting that changes in 3D chromatin structure could account, in part, for the conditional specificity of the eQTLs. P-values were calculated with a Wilcox test. (**H**) eQTL association plots colored by LD relative to FN-f lead variant rs10453229, a mega Hi-C heatmap, and ATAC-seq and RNA-seq signal tracks depicting an example of a distal FN-f-specific eQTL for *LPAR1* that is supported by a chromatin loop.

### Shared genetic architecture between eQTLs and GWAS variants reveals novel putative OA risk genes

To determine if any of our identified eQTLs could explain OA risk loci, we performed colocalization analysis between our eQTLs and 100 independent OA GWAS loci described by Boer et al.^2^ who mapped risk variants for 11 OA-related phenotypes including finger OA, thumb OA, hand OA, total hip replacement (THR), hip OA, all OA, knee-hip OA, knee OA, spine OA, total joint replacement (TJR), and total knee replacement (TKR). We identified 14 colocalized signals corresponding to 13 unique eGenes covering 6 different OA phenotype subtypes (**Table 1, Table S13;** coloc^46^; posterior probability (PP4) > 0.7). We identified 3 of the 5 previously reported chondrocyte e/pQTL/OA GWAS colocalized genes and added 10 novel colocalized eQTL signals. Only 1 of these colocalizations was identified as an eQTL in both conditions with 69.2% (9 of 13) and 23.1% (3 of 13) detected in only PBS or FN-f treated conditions, respectively. We performed colocalization analysis for all eQTL signals in both conditions regardless of whether or not they were detected as an eQTL in that condition. For several signals we observed colocalization even if the eQTL analysis did not meet our cutoffs for statistical significance (see *TGFA* below). Examples of colocalizations that were shared, PBS-specific, or FN-f-specific are highlighted in **Figure 5A-C**. The risk allele for the OA GWAS variant rs3771501 was associated with decreased expression of *TGFA* in PBS-treated chondrocytes (**Fig S8A**) and while the trend appeared the same in FN-f treated cells, the adjusted p-value (q-value = 0.057) did not reach our cutoff for statistical significance. Nevertheless it was identified as a colocalized signal in both conditions. In contrast, the risk alleles of the OA GWAS variants rs56132153 and rs9396861 were associated with decreased expression of *PIK3R1* and *RNF144B*, respectively, in their specific conditions (**Fig S8B-C**). These results underscore the importance of mapping eQTLs in a disease-relevant condition as well as a matched control for assessing risk before disease onset.

**Table 1.** Colocalizations between PBS and FN-f eQTL and OA GWAS signals. OA, osteoarthritis; GWAS, Genome-wide association study; THR, Total Hip Replacement; PP4, value of posterior probability 4 from coloc in the associated eQTL condition.

| gene | GWAS top OA phenotype | GWAS lead SNP | risk allele | risk allele frequency | risk allele association with expression | eQTL condition | PP4 | novel coloc-alization | eQTL genetic and condition interaction | change in OA | change with FN-f | change with age | M vs F |
| --- | --- | --- | --- | --- | --- | --- | --- | --- | --- | --- | --- | --- | --- |
| *ABCA10* | THR | rs2716212 | G | 0.3836 | higher | PBS | 0.888 | yes | - | - | - | - | - |
| *ABCA5* | THR | rs2716212 | G | 0.3836 | higher | PBS | 0.963 | yes | Both | down | - | - | - |
| *ABCA9* | THR | rs2716212 | G | 0.3836 | higher | PBS | 0.876 | yes | - | - | - | - | - |
| *ALDH1A2* | Hand OA | rs11071366 | T | 0.3865 | lower | PBS | 0.965 | no | - | up | - | - | - |
| *CARF* | All OA | rs62182810 | A | 0.5441 | higher | PBS | 0.778 | yes | - | - | down | - | - |
| *MAP2K6* | THR | rs2716212 | G | 0.3836 | higher | PBS | 0.983 | yes | - | - | down | - | - |
| *PAPPA* | THR | rs1321917 | C | 0.4088 | higher | Both | 0.963/0.881 | yes | - | up | up | up | - |
| *PIK3R1* | THR | rs56132153 | A | 0.6056 | lower | PBS | 0.903 | yes | PBS | - | down | up | - |
| *RNF144B* | Finger OA | rs9396861 | A | 0.6097 | lower | FN-f | 0.985 | yes | FN-f | down | up | - | - |
| *SLC44A2* | All OA | rs10405617 | A | 0.3194 | higher | FN-f | 0.954 | no | - | down | down | - | - |
| *SMAD3* | Hip OA | rs12908498 | C | 0.5384 | lower | FN-f | 0.98 | no | FN-f | - | - | - | - |
| *TGFA* | All OA | rs3771501 | A | 0.4679 | lower | PBS | 0.978 | yes | - | - | - | - | - |
| *TRIOBP* | THR | rs12160491 | G | 0.2887 | lower | PBS | 0.965 | yes | PBS | - | - | - | - |

**Fig 5:**
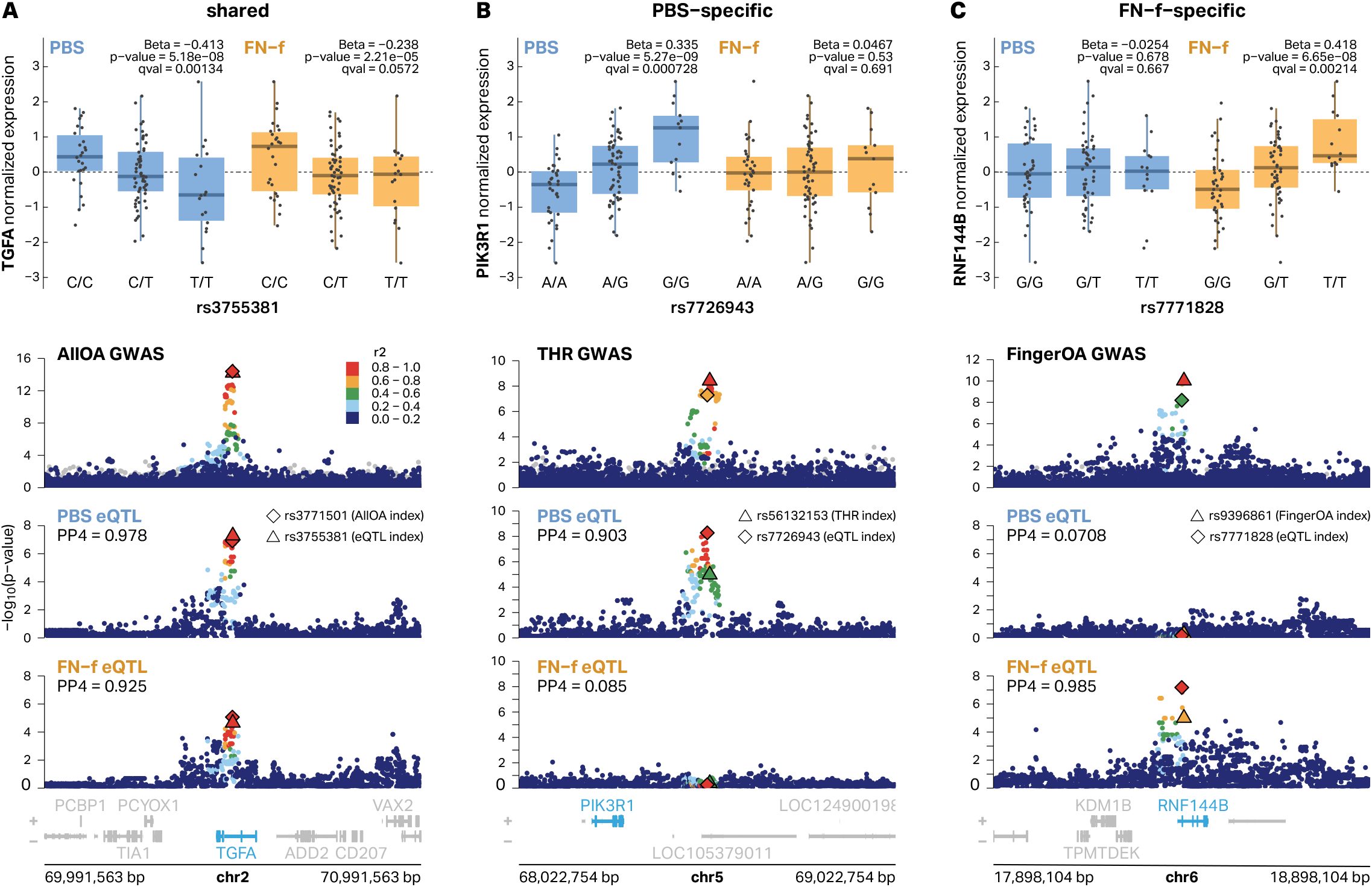
Examples of PBS-specific, FN-f-specific, and shared eQTLs that colocalize with OA GWAS signals. (**A**) eQTLs for TGFA are shared in both conditions and colocalize with an All OA GWAS signal with index variant rs3771501. eQTL signals are colored by LD relative to PBS lead variant rs3755381. (**B**) A distal eQTL signal for *PIK3R1* is specific to PBS-treated chondrocytes and colocalized with a Total Hip Replacement (THR) GWAS signal with index variant rs56132153. eQTL signals are colored by LD relative to PBS lead variant rs7726943. (**C**) An FN-f-specific eQTL signal for *RNF144B* colocalizes with a Finger OA GWAS signal with index variant rs9396861. eQTL signals are colored by LD relative to the FN-f lead variant rs7771828. All eQTL boxplots depict donor genotypes vs normalized eGene expression. All GWAS signals are colored by LD relative to the lead GWAS variant according to the 1000 Genomes European reference panel.

Many of the colocalized eGenes exhibited multiple lines of evidence linking them to a role in OA. Of the 13 colocalized eGenes (**Table 1**), 6 exhibited differential expression in response to FN-f (2 up and 4 down), 2 exhibited increased expression with age, and 5 were previously shown to exhibit expression changes in OA tissue^43,47,48^(2 up and 3 down). None of them exhibited sex-biased gene expression. Gene ontology and pathway enrichment analysis did not reveal any significant GO terms or pathways enriched in our set of 13 eGenes, which could be due to the low power associated with small (i.e. 13) sets of genes or could suggest that these genes influence OA risk through multiple distinct processes. Indeed the proteins coded by *ABCA10, ABCA5*, and *ABCA9* are all members of the ATP-binding cassette (ABC) transporter family, and *TGFA* and *SMAD3* encode proteins that regulate gene transcription and cellular proliferation.

One of the colocalized eGenes with multiple lines of support is the metalloproteinase pappalysin 1 (*PAPPA*). *PAPPA* was previously found to be upregulated in OA tissue^47^ and here we show that *PAPPA* exhibits increased expression in FN-f-treated cells (**Fig 6A**) and in older donors (**Fig 6B**). The lead GWAS risk variant (rs1321917) was identified with an odds ratio of 1.10 and is associated with increased expression of *PAPPA* in our eQTL analysis (**Fig 6C**). This locus was identified as an eQTL in both PBS and FN-f-treated chondrocytes and was colocalized with the OA GWAS signal for Total Hip Replacement (THR) in both conditions as well (**Fig 6D, Fig S9**). None of the GWAS variants at this locus (r^2^ > 0.6) overlap the promoter or gene body of *PAPPA* and the locus was assigned to *ASTN2* based on the nearest gene approach in Boer et al^2^. The lead GWAS variant is 409 Kb downstream of the *PAPPA* promoter and even the closest GWAS variant (LD r^2^ > 0.6) is 351 Kb downstream of the *PAPPA* promoter. However, a chromatin loop connects the promoter of *PAP-PA* to lead variants at the GWAS locus, which provides a possible mechanistic basis for this long-range regulation. This loop was recently described by Bittner et al., who provided further support for long-range communication at this locus by demonstrating that this GWAS signal colocalizes with a methylation QTL for a methylation site near the *PAPPA* promoter^49,50^. The *PAPPA* locus provides a model example of how a multi-omic approach can provide insight into the putative genes and mechanisms responsible for the contributions of particular genetic regions to OA risk.

**Fig 6:**
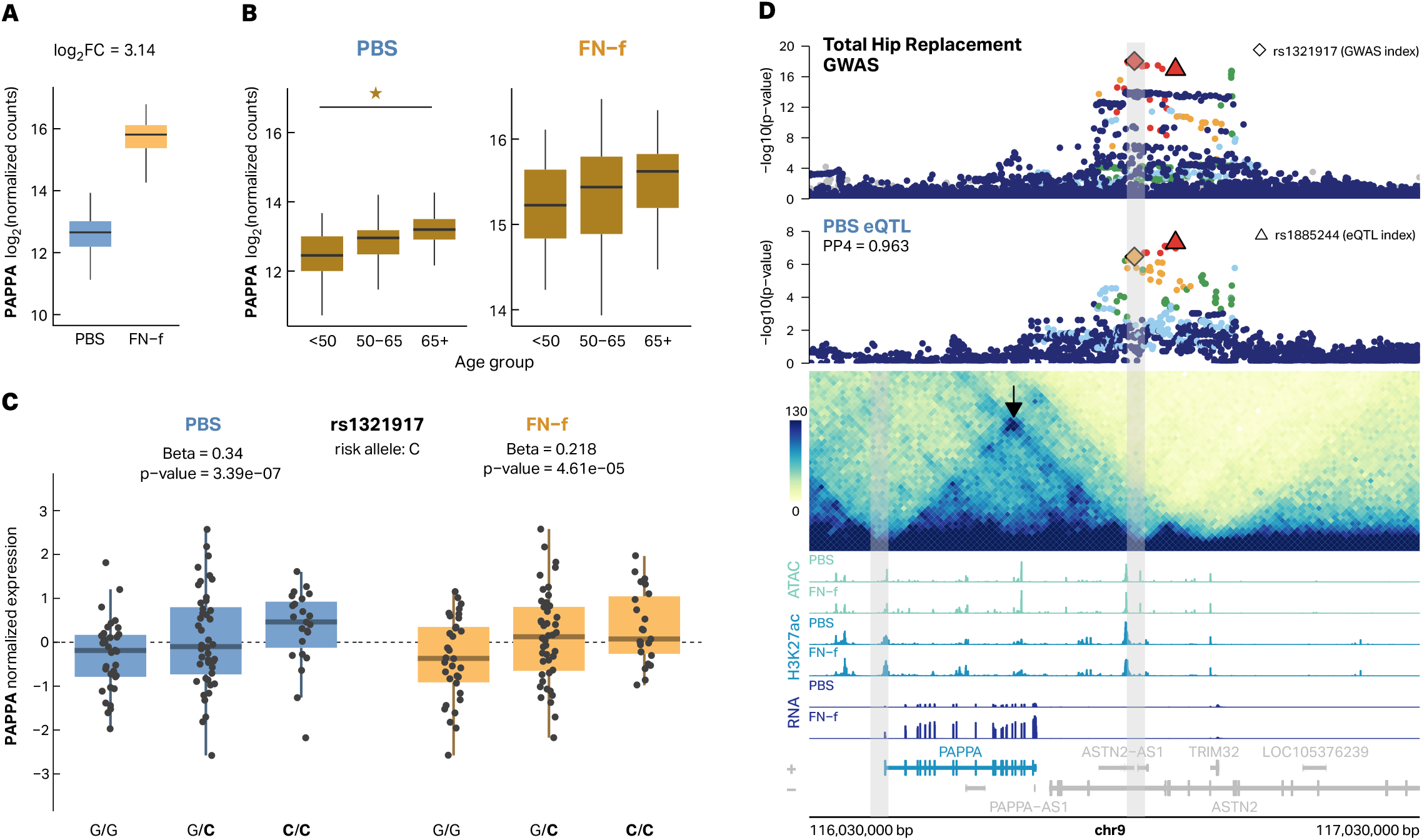
Multiple lines of evidence support *PAPPA* as an OA risk gene. The matrix metalloproteinase *PAPPA* has previously been found to have increased expression in OA tissue. (**A**) A boxplot depicting increased *PAPPA* log2 normalized expression counts in response to FN-f treatment (log2 fold change = 3.14). (**B**) *PAPPA* shows increased expression in older donors. Star indicates significance (adjusted p-value < 0.05). (**C**) The OA risk allele (C, bolded) of variant rs1321917 is associated with higher expression of *PAPPA* in chondrocytes in both PBS (blue) and FN-f (yellow). (**D**) The colocalized PBS eQTL signal for *PAPPA* does not overlap with the promoter or the gene body but is connected to the promoters via a 420 Kb chromatin loop which supports this distal eQTL. Total Hip Replacement GWAS is colored by LD relative to the GWAS index rs1321917 according to the 1000 Genomes European reference panel. Gray bars highlight the genomic regions of loop anchors.

## Discussion

By mapping expression, chromatin accessibility, and 3D chromatin structure across individuals and conditions we provided critical new insights into the mechanisms driving genetic risk for OA. We identified thousands of genes that are differentially expressed in chondrocytes responding to FN-f, an OA-related stimulus that models cartilage matrix damage. These expression changes correlated with those seen in OA tissue supporting the use of this system to study OA-related chondrocyte gene regulation. We provided comprehensive, highly-powered characterizations of human chondrocyte gene expression differences related to age and sex, two important risk factors for OA. We then mapped eQTLs and response eQTLs to reveal how common genetic variation contributes to chondrocyte gene expression both in resting and activated conditions. We mapped changes in 3D chromatin architecture in chondrocytes responding to FN-f and found gained loops and increased expression at many key OA risk genes including *JUN, NFKB*, and *MMP13*. Many of these loops and changes in chromatin structure provided mechanistic insight and explanation for our distal and condition-specific eQTLs. Finally, we colocalized our eQTLs with 11 OA GWAS phenotypes revealing 13 putative OA risk genes, 76.9 percent of which have not been described by previous chondrocyte QTL studies.

Our eQTLs included 4 of the 5 e/pQTLs previously colocalized to OA GWAS signals identified from UK Biobank data^3^. This included three of the four previously colocalized^16^ chondrocyte eQTLs. Two of those (*SMAD3* and *SLC44A2*) were also colocalized with OA GWAS variants in our analysis. The third eQTL (*NPC1*) was not identified as colocalized in our analysis but only because that locus was no longer significant in the updated OA GWAS study that we used (**Fig S10A**). Our colocalized eQTLs also identified a colocalized eQTL that was previously identified^16^ as a colocalized pQTL (*ALDH1A2*). Only one of the previously identified colocalized QTLs (*FAM53A*) was not identified in our analyses (**Fig S10B**), which we suspect is due to differences between our study designs, where we used healthy tissue treated with FN-f and the previous study used primary OA tissue.

Characterization of our 13 colocalized eGenes both independently and collectively provides important insights into the genetic basis of OA and possible strategies for therapeutic development. Some of these genes have known roles in OA or are involved in processes relevant to OA pathology while the function of others is less clear. SMAD3 is a key transcriptional regulator in the TGF-B signaling pathway and SMAD3 disruption has been shown to cause an OA-like phenotype in mouse models^39^. ALDH1A2 is involved in retinoic acid synthesis and low ALDH1A2 in chondrocytes has been previously associated with increased expression of inflammatory genes that are also un-regulated in response to articular cartilage injury^51^. SLC44A2 is a choline transporter whose role in OA is not well understood.

Importantly, this study revealed 10 novel eGenes (*ABCA5, 9*, and *10, CARF, MAP2K6, PAPPA, PIK3R1, RNF144B, TGFA*, and *TRIOBP*) that colocalized with OA GWAS signals, providing new insights into the etiology of OA. *ABCA5, 9*, and *10* are genes that cluster on chromosome 17q24.3 and code for members of the ATP binding cassette (ABC) subfamily, a group of proteins that serve to transport a variety of molecules across membranes^52^. A role for this transporter family in cartilage biology or OA has not been investigated. *CARF* codes for a calcium responsive transcription factor which also has not been previously investigated in the context of OA. MAP2K6, also known as MEK6, is a member of the MAP kinase signaling family and phosphorylates p38, which is a mediator of catabolic signaling in cartilage that includes signaling activated by FN-f and cytokines such as IL-1^53^. Inhibition of p38 in vitro can inhibit cartilage degradation^53^ although genetic inhibition of p38 in transgenic mice expressing a dominant negative p38 construct resulted in more severe OA at 1 year of age^54^. PIK3R1 is a regulatory subunit of the PI-3 kinase. The role of PI-3 kinase signaling in cartilage biology and OA is complex. PI-3 kinase is a positive mediator of chondrocyte anabolic activity and cell survival but is also activated by pro-inflammatory cytokines including IL-1 and oncostatin M which promote catabolic signaling^55^. *RNF144B* codes for a ring finger protein that regulates ubiquitin-protein transferase activity. It can inhibit LPS-induced inflammation^56^ which may be relevant to inflammation in OA^57^. The role of TGFα and its activation of the EGF receptor in OA is also complex. TGFα has been shown to induce chondrocytes to produce catabolic factors such as MMP13^58^, but in contrast the intra-articular injection of nanoparticles delivering TGFα reduced the severity of surgically-induced OA in mice^59^. TRIOBP is the TRIO and F-actin binding protein which stabilizes F-actin structure TRIOBP was recently found to co-localize with eQTLs from human osteoclast-like cells generated from isolated human peripheral blood mono-nuclear cells^60^ but its role in cartilage biology and OA has not been studied.

Among the novel colocalized eGenes, *PAPPA* is particularly interesting. *PAPPA* is a zinc metalloproteinase that cleaves IGF binding proteins including IGFP-4 and -5^61^. It exhibits increased expression in OA tissue, in older donors, and in response to FN-f. The shared GWAS and eQTL signal is over 400 Kb away from the promoter of *PAPPA* but as we (and others^49^) have shown, these variants are connected to the promoter of *PAPPA* via a chromatin loop. A recent study pinpointed *PAPPA* as the most consistent mediator of senescence induction in sirtuin-deficient human induced pluripotent stem cells^62^. Of note, the authors identified a loop with an enhancer locus 416 Kb downstream of the *PAPPA* promoter as emerging in response to knockout of sirtuins 1 and 5 (a sirtuin deficiency-sensitive genomic region), which overlaps the eQTL/GWAS region we identified in chondrocytes. *In vitro* and *in vivo* studies support a role for *PAPPA* secretion in amplifying senescence^63^ and limiting median lifespan^64^. Given the importance of IGF signaling in cartilage homeostasis and repair^65^, as well as the potential role for senescence to drive joint dysfunction^66^, alterations in *PAPPA* expression or function could potentially influence the balance between anabolic and catabolic processes in the joint. Further functional studies are required to delineate *PAPPA*’s exact role in OA risk and whether modulation of *PAPPA* has any therapeutic potential.

Interestingly, the identified genes are not clearly enriched in any specific pathways or biological processes which could indicate a number of different things. First, the GWAS data describes 11 different phenotypes ranging from finger OA to total hip replacement. Different OA subtypes and phenotypes measured by these GWAS may be driven by distinct mechanisms and involve distinct biological pathways. Second, OA is a polygenic disease influenced by a number of environmental factors. It is possible, even likely, that OA risk is driven by genetic influences on a number of different pathways and processes that act across a diverse array of developmental time points and biological conditions. As such, one would not expect to see enriched ontologies or pathways until a larger set of risk genes was identified.

Despite the success of this project it is important to highlight several limitations. First, while this study successfully identified 13 putative OA risk genes, eQTLs alone are not well suited to pinpointing the exact causal variants at each locus. Other genomic methodologies including chromatin accessibility QTL mapping and massively parallel reporter assays could be useful in defining causal variants. Second, while we more than doubled the number of colocalized eQTLs and OA risk variants, many OA GWAS loci remain unexplained. Increasing our sample size will increase our power and likely lead to more colocalized eGenes (particularly those controlled by distal regulatory elements). While much of the genetic contribution to OA risk is thought to be mediated via chondrocyte function, some variants surely impact other cell types and other components of the joint. Mapping eQTLs in additional cell types including fibroblasts, macrophages, and other cell types present in the joint will likely increase the number of colocalized signals. Moreover, while many of the variants are likely to impact resting or FN-f-activated chondrocytes, some may impact chondrocytes during development or in response to other stimuli. Finding ways to interrogate the genetic influence on gene expression across other conditions may also reveal putative risk genes. Finally, while colocalized eQTLs provide strong support for causal associations, final proof of this causal effect and assessment of therapeutic potential will require focused functional studies.

This study represents a major breakthrough for OA research by providing potential explanations for 13 OA GWAS risk loci. The next critical steps are to functionally characterize the role that these genes play in the OA phenotype and determine if modulating their expression or activity can alleviate or even reverse OA-related symptoms. In parallel, it will be important to continue generating similar data sets across multiple cell types and conditions with increased sample sizes to improve our power to detect OA risk genes.

## Methods

### Sample collection and treatment

Chondrocytes from human talar cartilage of deceased tissue donors with no known history of arthritis (see **Table S14** for donor characteristics) were isolated via enzymatic digestion and cultured in monolayer using protocols as previously described^19^. Chondrocytes were given 4 days of recovery from digestion and maintained in cell culture medium consisting of DMEM/F12 supplemented with 10% FBS (VWR Seradigm; #97068-085) before experiments. For DNA experiments, cultured chondrocytes were first collected as cell pellets via trypsinization and stored at -80° until DNA isolation. For FN-f treatment and RNA isolation, serum-containing media was first removed, and the cells washed twice with PBS before being serum-starved in DMEM/F12 for 2 hrs. Chondrocytes were then treated with 1 μM purified recombinant human FN-f (FN7-10), prepared as previously described and stored as aliquots at -80 degrees C in PBS^23^, or treated with PBS alone as a control for 18 hours. The media was then removed, and the cells washed once with PBS before lysis using RNeasy Lysis Buffer (Qiagen). Lysates were stored at -80° until RNA isolation and purification.

### DNA extraction

Genomic DNA was extracted from chondrocytes using the QIAamp DNA mini kit (Qiagen, #51304) according to the manufacturer’s instructions. Samples were quantified with Qubit High Sensitivity assay kit (Thermo Fisher Scientific #Q32854) and absorbance values were obtained using NanoDrop. DNA was submitted to the Mammalian Genotyping Core at University of North Carolina to be genotyped using the Infinium Global Diversity Array-8 v.10 Kit (Illumina #20031669).

### Genotype processing and quality control

SNP genotypes were exported into PLINK format with the Illumina software GenomeStudio. Quality control and filtering was performed with PLINK (v1.90b3.45)^67^. We filtered out SNPs with missing genotype rate > 10% (--geno 0.1), deviations from Hardy-Weinberg equilibrium at a p-value < 1 x 10e-6 (--hwe 10^-6), and minor allele frequency < 1% (--maf 0.01). Samples with sex discrepancies from PLINK –check-sex comparison between reported sample sex an sex assigned from heterozygosity on the X chromosome were omitted. To assess relatedness of samples, identity by descent (IBD) was calculated with PLINK. Samples were retained if their inferred relationship type was either UN (unrelated) or OT (other) with PI_HAT (proportion IBD) < 0.2. To estimate the population structure of our samples, we combined our data with overlapping data from the 1000 Genomes Project^68^ and used EIGENSTRAT (v8.0.0)^69^ to conduct principal component analysis (PCA) optimized for population-related analyses. Prior to imputation, we filtered our dataset for autosomes, flipped the alleles of SNPs that were not on the reference strand as identified by snpflip^70^, and converted PLINK files into VCF files separated by chromosome. Data was imputed using the version R2 on GRC38 TOPMed reference panel with Eagle2 (v2.4) phasing^71^ on the TOPMed Imputation Server^72^. Following imputation, we followed similar QC filtering steps as before imputation and retained SNPs with missing genotype rate < 10%, p-value of Hardy-Weinberg equilibrium > 1 x 10e-6, minor allele frequency > 1%, and sufficient imputation quality (R2 > 0.3). The resulting final dataset contained approximately 9.7 million autosomal SNPs.

### RNA isolation

RNA was extracted using the RNeasy kit (Qiagen #74104) according to the manufacturer’s recommendation. On-column DNase digestion was performed during the extraction. Samples were quantified with the Qubit RNA High sensitivity assay kit (Thermo Fisher Scientific #Q32582) and RNA integrity number (RIN) was obtained using the Agilent TapeStation 4150. RNA was submitted to the New York Genome Center for RNA-seq library preparation and sequencing.

### RNA-seq processing and quality control

RNA-seq libraries were sequenced at the New York Genome Center to an average read depth of approximately 101 million paired end reads (2 x 100 bp) per sample. FASTQ files sequenced on multiple flow cells but were from the same library were merged. After trimming low quality reads and adapters with TrimGalore! (v0.6.7)^73^, we performed quality control of each library with FastQC (v0.11.9)^74^. Trimmed FASTQs were aligned against the GENCODE.GRCh38.p13 reference genome with STAR aligner (v2.7.10a)^75^ and obtained transcript-level quantifications with salmon (v1.10.0)^76^ with –gcBias and –seqBias flags and the ENSEMBL version 97 (GRCh38.p12) hg38 cDNA assembly. To conduct differential gene expression analysis, transcript-level quantifications for each sample were summarized and converted to gene-level scaled transcripts in R with tximeta^77^. Individual donor RNA signal tracks were created with deepTools (v3.5.1)^78^ and then merged by condition.

Evaluation of sample swaps and sample contamination was performed with VerifyBamID (v1.1.3)^79^. Genotyping sample swaps (n = 2) were corrected. Samples with FREEMIX and CHIPMIX scores > 0.2 after attempting to fix sample swaps were omitted.

MultiQC (v1.11)^80^ aggregated QC results from FastQC, STAR, salmon, and VerifyBamID. Samples with > 10% unmapped short reads, samples without a corresponding QC’d genotyping sample (see below), and donors without both a PBS RNA-seq sample and FN-f RNA-seq sample that passed QC were omitted. By all these criteria, the final datasets included 101 individual donors, corresponding to 202 RNAseq samples (101 PBS and 101 FN-f).

### Replicate correlation

Technical replicates (n=2 for PBS and n=3 for FN-f) were performed for the RNA-seq analysis using chondrocytes cultured from three donors. For each treatment, VST-normalized gene expression counts were used to calculate Pearson’s correlations between libraries from the same donors and between libraries across different donors. Correlation coefficients were transformed with Fisher’s z. Significance of difference between donor-self libraries and donor-other libraries was tested with an unpaired, two-sided Wilcox test.

### Differential analysis of FN-f-induced transcriptional changes

Differential analysis between FN-f samples and PBS samples was conducted in R with DESeq2^24^ using summarized gene-level scaled transcripts. A design of ∼Donor + Condition was used to adjust for donor variability while calculating changes between PBS and FN-f conditions. Before modeling, lowly expressed genes were omitted by requiring at least 10 counts in 10% of samples. Shrunken log2 fold change values were calculated using the “apeglm” method of lfcShrink^81^. Genes were considered differential with an FDR-adjusted p-value < 0.05 (Wald test) and shrunken absolute log2 fold change > 1. These genes were further filtered for the largest and most significant threshold using an FDR-adjusted p-value < 0.01 and absolute log2 fold change > 2.

### GO term, KEGG pathway, and transcription factor motif enrichment of differential FN-f genes

Filtered high-significance differential FN-f genes (padj < 0.01 and absolute log2 fold change > 2) were split based on direction of effect. findMotifs.pl in the HOMER software suite (v4.11)^25^ was used on these groups to identify significantly enriched GO Terms (p < 0.01), KEGG pathways (p < 0.01), and transcription factor motifs (p < 0.01). GO terms were reduced based on semantic similarity using rrvgo (v1.14.2)^82^.

### Comparison to publicly available OA gene expression datasets

Differential expression data in OA tissue was used from 3 published studies as a comparison to our datasets. Microarray gene expression results between OA and preserved cartilage were downloaded from the RAAK study^47^ and filtered for genes with p-value < 0.05. The list of all genes detected in an RNA-seq analysis comparing normal and OA knee cartilage was obtained from Fisch et al. (2018)^43^ and filtered for genes with an adjusted p-value < 0.05. Since a supplementary list of differential gene expression results was not readily accessible, the non-normalized count matrix from Fu et al. (2021)^48^ was downloaded from GEO under accession number GSE168505 and analyzed with DESeq2^24^. Genes with at least 10 counts in 1 sample were included in the analysis. Since no additional covariate information was available, differential expression between OA and normal cartilage was tested using a design of ∼Condition. Shrunken log2 fold change values were calculated using the “apeglm” method of lfcShrink^81^ and results were filtered for differential genes with an FDR-adjusted p-value < 0.05. A final set of differential OA genes was defined as genes that were significant and showed the same direction of effect in all 3 studies. A Wilcox test was used to determine if the FN-f induced log2 fold change of upregulated and downregulated OA genes were significantly higher or lower, respectively, than genes not found in this set.

### Sex-specific gene expression analysis

Summarized gene-level transcript results were separated based on condition and analyzed for sex-specific effects using DESeq2^24^. A design of ∼Ancestry + Age_group + Sex was used to control for donor genetic ancestry (as determined from principal component analysis with 1000 Genomes samples using EIGENSTRAT with 1000 Genomes-defined superpopulations; AFR, AMR, EAS, EUR, or SAS) and donor age group (31-40, 41-50, 51-60, 61-70, 71-80, or 81-90) while assessing differences in sex-related expression. lfcShrink^81^ was used to calculate shrunken log2 fold change values using the “apeglm” method. Genes were considered significantly sex-specific with an FDR-adjusted p-value < 0.01. A union of sex-specific genes found in either PBS or FN-f samples was used for downstream analyses. Sex-specific genes were considered differentially expressed in OA tissue if the gene was significant (adjusted p-value < 0.05) in any of the 3 OA studies described above.

### Comparison of sex-specific genes to GTEx sex-biased gene expression

To compare human chondrocyte sex-specific gene expression to other sex-biased expression in other tissues, summary statistics of GTEx sex-biased genes in 44 tissues^30^ was downloaded from the GTEx portal (https://gtexportal.org/home/datasets). Datasets were compared based on ENSEMBL gene ID.

### Identifying genes with age-dependent expression patterns

DESeq2^24^ was used to identify genes with age-related expression patterns in summarized gene-level transcripts separated by condition. A likelihood ratio test (LRT) was used to test dependence of counts on a smooth function of age, by modeling age with natural cubic splines with five degrees of freedom^83^. To control for donor sex and donor genetic ancestry, the full model was ∼Sex + Ancestry + splines::ns(Age, df = 5) and the reduced model was ∼Sex + Ancestry where Ancestry was determined from principal component analysis with 1000 Genomes samples using EI-GENSTRAT with 1000 Genomes-defined superpopulations (AFR, AMR, EAS, EUR, or SAS). Genes were considered significantly age-related if the adjusted p-value of the LRT was < 0.05. k-means clustering of centered fitted spline curves with a k of 2 was used to assign to gene clusters exhibiting increased expression with age and decreased expression with age. GO term enrichment for each of these clusters was performed using findMotifs.pl in the HOMER software suite (v4.11)^25^. GO terms were reduced based on semantic similarity using rrvgo (v1.14.2)^82^ and considered significant with p < 0.01. Age-related genes were considered differentially expressed in OA tissue if the gene was significant (adjusted p-value < 0.05) in any of the 3 OA studies described above.

### Comparison of age-related genes to GTEx age-related gene expression change

To compare age-related genes in human chondrocytes to other tissues, aging-related statistics for genes in nine human tissues was downloaded from Yang et al. (2015)^33^. For consistency with Yang et al., Thyroid and Skin tissues were omitted from the dataset and genes were considered significantly age-associated with an FDR-adjusted p-value < 0.05. Datasets were compared based on ENSEMBL gene ID.

### ATAC-seq library preparation

Chondrocytes were treated with FN-f or PBS for 18 hours as described above, media was aspirated and cells were washed with PBS. To avoid changes associated with trypsinization, cells were directly lysed in the well as previously described^84^. Briefly, cells were washed twice with cold PBS followed by one wash with cold ATAC-seq resuspension buffer (RSB). Cells were lysed in RSB containing 0.1% NP40, 0.1% Tween-20, and 0.01% digitonin for 10 min at 4C. After lysis the remainder of the Omni-ATAC protocol was performed^85^. Following washes and transposition with Tagment DNA TDE1 Enzyme (Illumina #20034197) reactions were cleaned up with DNA clean and concentrator kit (Zymo Research #D4014). Samples were preamplified using High-Fidelity 2X PCR Master Mix (New England Biolabs, #M0541L) and adapters (Illumina Nextera XT Index kit #FC-131-1001). The number of additional cycles was determined by quantitative PCR. Following a double-sided AMPure XP bead cleanup (Beckman Coulter #A63881), libraries were quantified using Qubit. Library quality and fragment distribution was visualized by Agilent TapeStation 4150. Prior to pooling, libraries were quantified with the KAPA library quantification kit (Roche #07960298001). Libraries were sequenced on Illumina NextSeq 500 sequencer (75-bp paired-end reads, high output kit Illumina #20022907) at the CRISPR core, University of North Carolina.

### ATAC-seq data processing

Adaptors and low-quality paired-end reads were processed using Trim Galore! (v0.6.7)^73^. Reads were then aligned to the UCSC hg38 human genome reference using BWA-MEM (v0.7.17)^86^. We removed duplicate alignments with Picard (v2.10.3)^87^ and excluded mitochondrial reads via samtools (v1.17)^88^. Quality assessment of ATAC-seq data, including total read counts, duplicate rates, transcript start site enrichment scores, and the fraction of reads in called peak regions, was conducted using R package ATACseqQC (v3.18)^89^. All samples met the ENCODE project’s standards as of July 2020. We eliminated reads mapping to ENCODE blacklist regions (Accession ID: ENCFF356LFX) using bedtools (v2.30)^90^. To adjust for the Tn5 transposase binding bias, we applied a Tn5 shift correction with alignmentSieve from deepTools (v3.5.1)^78^. Peak calling was performed with MACS3 (v3.0.0)^91^, utilizing the following parameters: ‘callpeak -f BAM --call-summits -B -q 0.01 --nomodel --shift -100 --extsize 200 --keep-dup all’. We merged peaks identified under two different conditions using bedtools (v2.3.0)^90^.

### Differential ATAC peak analysis, chromatin accessible region overlap with eQTLs, and transcription factor motif enrichment

To identify ATAC peaks that were differentially accessible between FN-f and PBS samples, we used DESeq2^24^ with peak read counts described above using a design of ∼Donor + Condition to adjust for donor variability. Prior to testing, we filtered for peaks with at least 10 counts in 2 samples. Shrunken log2 fold change values were calculated with the “apeglm” method of lfcShrink^81^. Peaks were considered differentially accessible with globally adjusted p-value < 0.01 and shrunken absolute log2 fold change > 1.

We overlapped all called peaks in either condition with high confidence PBS-specific lead eQTLs, shared lead eQTLs, and high confidence FN-f-response lead eQTLs and variants in high LD (r^2^ > 0.8) with these groups with findOverlaps from the GenomicRanges R package^92^. To test for enrichment of condition-specific accessibility of condition-specific eQTLs, we performed a Wilcox test comparing the peak log_2_ fold change values of peaks overlapping condition-specific eQTLs to peaks that overlapped any lead eQTL or variant in high LD (r^2^ > 0.8). An alternative hypothesis of “less” was used for testing peaks overlapping PBS-specific eQTLs and an alternative hypothesis of “greater” was used for testing peaks overlapping FN-f-specific eQTLs.

Transcription factor motif enrichment of peaks overlapping high confidence condition-specific PBS eQTLs and peaks overlapping high confidence FN-f-response eQTLs was performed using findMotifsGenome.pl in the HOMER software suite (v4.11)^25^. Enrichment was calculated against a background of any peak that overlapped any lead eQTL or variant in high LD (r^2^ > 0.8) in either condition.

### *In situ* Hi-C library preparation

4 donor plates of 8 million chondrocytes were cultured in DMEM/F-12 media, serum-starved for 2 hours, and treated with PBS or FN-f. After 18 hours of treatment, the media was removed from the plate. Cells in each plate were crosslinked in 10% formaldehyde in DMEM/F-12 media and incubated for 10 minutes on a rocker. To quench, 2M Glycine was added as a final concentration of 0.2M and incubated for 5 minutes on the rocker. The supernatant was removed, the cells were resuspended with 10mL cold PBS, collected into a 15mL tube, and spun down at 2500 rpm, 4°C for 5 minutes. The pellets were resuspended with 1mL PBS, transferred to 1.5mL microcentrifuge tube, and spun down at 900g, 4°C for 5 minutes. The pellets (∼8 million cells) were flash frozen in liquid nitrogen and stored at -70 °C. The cells were thawed and *in situ* Hi-C was performed as described in Rao et al. (2014).^4^

### Hi-C data processing

Hi-C data was processed using the modified Juicer pipeline (https://github.com/EricSDavis/dietJuicer) with default parameters, as previously described^93^. Reads were aligned to the hg38 human reference genome with bwa, and MboI was used as the restriction enzyme. A total of 3,170,331,152 Hi-C read pairs were processed from PBS-treated chondrocyte cells, resulting in 1,949,761,524 Hi-C contacts (61.5%). Similarly, 2,925,877,690 Hi-C read pairs were processed from FNF-treated chondrocyte cells, yielding 1,836,062,944 Hi-C contacts (62.75%). Hi-C matrices were constructed individually for each of the two technical replicates across four biological replicates. Subsequently, the Hi-C mega map was merged with all replicates about each condition (PBS or FNF-treated chondrocytes).

Loops were identified at 5 kb resolution with Significant Interaction Peak (SIP) caller (v1.6.2)^94^ and Juicer tools (v2.13.07) using the replicate-merged mapq >30 filtered hic file with the following parameters: ‘-norm SCALE -g 2.0 -min 2.0 -max 2.0 -mat 2000 -d 6 -res 5000 -sat 0.01 -t 2000 -nbZero 6 -factor 1 -fdr 0.05 -del true -cpu 1 -isDroso false’.

### Differential loop analysis

DESeq2^24^ Wald testing was used for differential analysis of loops using a model of ∼Condition + Donor + replicate. Shrunken log2 fold change values were calculated with the “apeglm” method of lfcShrink^81^. Loops were considered differential with a globally adjusted p-value < 0.1. We identified protein-coding gene promoters that overlapped either anchor of differential or static loops using the GENCODE Release 44 hg38 (GRCh38.p14) reference genome and findOverlaps function^92^. GO term enrichment analysis of these genes at differentially gained loop anchors was conducted using findMotifs.pl in the HOMER software suite (v4.11)^25^ against a background of genes at static loop anchors.

### Contact frequency between distal eSNPs and eGenes

We considered the range of an eQTL signal to span the minimum and maximum range of variants in moderate LD (r^2^ > 0.6) with the index variant to maximize capturing the entire signal width and any plausible putative variants. We investigated long-range contacts between SNPs and their eGenes by defining distal eQTL signals as those with the minimum or maximum signal range at least 50 Kb away from either end of the entire eGene. Connections between signals and eGene promoters via a chromatin loop (differential or static) were identified using the linkOverlaps function from InteractionSet^95^ with loop anchors expanded to 30 Kb.

Contact frequency count data between lead SNPs and gene promoters according to hg38 were extracted from PBS and FN-f mega map Hi-C files at 5 Kb resolution with SCALE normalization with pullHicPixels from the mariner R package^96^. The matchRanges function from nullranges^97^ was used to generate a null distribution of distance-matched SNP-gene pairs for testing contact frequency between lead SNPs and their assigned eGenes. A Wilcox test was used to determine if the contact frequency between SNPs and their eGenes was higher compared to the contact frequency between distance-matched SNP-gene pairs.

Contact frequency count data between eGenes and high-confidence PBS-specific, shared, and high-confidence FN-f-specific eQTLs and variants in high LD (r2 > 0.8) were also extracted at 5 Kb resolution and SCALE normalization with pullHicPixels from mariner. A log2 fold change in contact frequency was calculated for these pixel counts between the FN-f and PBS conditions. A Wilcox test was used to test for enriched contact frequency of condition-specific eQTLs with their associated eGenes in their associated condition.

### Condition-stratified *cis* eQTL mapping

Prior to eQTL mapping, we filtered out lowly expressed genes and only considered protein-coding genes that had at least 10 counts in more than 5% of all samples (11 samples). Samples were normalized using the “TMM” method from edgeR^98^. Gene expression data was then normalized separated by condition with an inverse normal transformation across each gene. The transcription start site (TSS) of each gene was defined as the start of the most upstream transcript according to the GENCODE Release 44 hg38 (GRCh38. p14) genome build.

Genetic variants were selected for testing with at least 10 counts of the minor allele and at least 5 heterozygote donors using GATK VariantFiltration^99^. For each gene, we considered variants within a 1 Mb window in either direction of the defined TSS.

Principal component analysis was performed on genotyping data with QTLtools pca^100^. The kneedle algorithm^101^ was used to identify the “elbow” of principal components versus percent variance explained to determine the number of genotyping principal components to include as covariates in our linear model. To infer technical confounders, we applied probabilistic estimation of expression residuals (PEER)^102^ to the condition-separated inverse-normalized gene expression results. To identify the number of PEER factors to include as covariate in each model, we generated PEER factors from 1-50 and performed QTL mapping with QTLtools (v1.3.1)^100^. A permutation-based analysis was performed with the QTLtools cis permutation pass with 1000 permutations. Adjusted empirical p-values were adjusted globally using the Storey-Tibshirani q-value^37^. eGenes, or genes with at least one significant eQTL, were defined with a q-value < 0.05 (an equivalent p-value of 8.64e-24 in PBS and 5.95e-21 in FN-f). We selected the final number of PEER factors to include in the model that yielded the most significant eGenes before a plateau in the number of significant eGenes with a successive increase in PEER factors. The final eQTL model for PBS samples was *expression ∼ SNP* + 4 *genotyping PCs* + 20 *PEER factors* + *Donor Sex* and the final eQTL model for FN-f samples was *expression ∼ SNP* + 4 *genotyping PCs* + 22 *PEER factors* + *Donor Sex*. eQTL nominal p-values were calculated with the QTLtools cis nominal pass. For each eGene, we obtained the local nominal threshold by calculating a p-value as the mean of the smallest p-value above the q-value threshold and the highest p-value above the q-value threshold and using the beta distribution (qbeta) with shape1 and shape2 parameters defined from the QTLtools permutation analysis, as described by FastQTL^103^. PBS eGene nominal thresholds ranged from 5.03e-6 to 3.39e-4 and FN-f eGene nominal thresholds ranged from 5.96e-6 to 4.19e-4.

To identify independent signals for each significant eGene, we performed conditional analysis with the QTLtools cis conditional pass using the above eGene nominal p-value thresholds and same set of covariates as the original eQTL models. rsIDs for independent variants were assigned based on position and allele-matching relative to the GRCh38.p14 build 156 dbSNP reference. After isolating conditionally distinct lead eQTL-eGene pairs, conditional signals for eGenes with more than 1 independent signal were isolated by re-running QTLtools cis nominal pass and conditioning on the lead variant(s) of the eGene’s other distinct signal(s).

### Comparison to existing cartilage eQTLs

High-grade and low-grade cartilage eQTLs from Steinberg et al. (2021)^16^ were downloaded from the Musculoskeletal Knowledge Portal (https://msk.hugeamp.org/) and were lifted over to hg38 with UCSC liftOver^104^ for compatibility with our dataset. To determine effect sizes of shared eGenes, we used the lead variant identified by Steinberg et al. The beta values of shared variants were adjusted so they were all in reference to the minor allele.

### Condition-specific and response eQTLs

Condition-specific and response eQTLs were identified by testing significant (q-value < 0.05) PBS and FN-f eGenes for the significance of an interaction term between genotype and condition. The R package lme4^105^ was used to compare the following two linear mixed models for all lead eSNP-eGene pairs: *H*0: *expression ∼ SNP* + *covariates* + *condition* + (1|*Donor*) *H*1: *expression ∼ SNP* + *covariates* + *SNP:condition* + (1|*Donor*) where covariates are the same covariates used in standard eQTL mapping, condition = 0 or 1 (PBS or FN-f, respectively), and (1|Donor) accounts for any donor-specific random effects. Interaction p-values were calculated using ANOVA. eQTLs with an interaction p-value < 0.05 were considered significant. We further filtered this list for a set of high-confidence PBS-specific and FN-f-response eQTLs by filtering for eQTLs that were only found in one condition, had at least 5 donors with each variant genotype, and had a beta difference of at least 0.2 between conditions. KEGG pathway enrichment for these eGenes was performed using findMotifs.pl in the HOMER software suite (v4.11)^25^

### Colocalization between eQTLs and OA GWAS

To test for colocalization between independent eQTL signals and OA GWAS, we used summary statistics for 11 OA phenotypes from Boer et al. (2021)^2^. Data was downloaded from the Musculoskeletal Knowledge Portal (https://msk.hugeamp.org/) and lifted over to hg38 coordinates with UCSC liftOver^104^. LD proxies (r^2^ > 0.8) of 100 lead variants (omitting sex-specific and early-onset OA phenotypes) were identified using the 1000 Genomes European reference panel since 11 of 13 GWAS cohorts were of European descent. PLINK (v1.90b3.45)^67^ –ld was used to calculate r2 values with the following parameters: –ld-window 200000 –ld-window-kb 1000.

We performed colocalization analysis between an eQTL and GWAS signals if the lead eQTL variant was in moderate LD (r^2^ > 0.5) with the lead GWAS variant according to either our in-study reference panel or the 1000 Genomes European reference panel. For each analysis, we considered the index GWAS variant and any variants within ± 250 Kb and filtered eQTL data for this same set of variants. We ran coloc.abf^46^ using default priors with eQTL data inputs of nominal p-values, sample size, minor allele frequencies, betas, and beta variances and GWAS data inputs of nominal p-values, minor allele frequencies, and betas. We considered a coloc posterior probability (PP4) > 0.7 as sufficient evidence of colocalization.

## Visualization

Gene expression heatmaps for condition, sex, and age were made using ComplexHeatmap^106^. Association plots, Hi-C maps, and other genomic signal tracks were plotted with plotgardener^107^. All other plot types were made with ggplot2^108^.

## Data Availability

The genotyping, RNA-seq, ATAC-seq, and Hi-C data sets generated for this research study are in the process of being submitted to the NIH’s database of Genotypes and Phenotypes (dbGaP) under the accession number phs003581. v1.p1. Full eQTL summary statistics are available through the Downloads page on the Musculoskeletal Knowledge Portal (https://msk.hugeamp.org/downloads.html).

## Acknowledgments

We thank Jason Stein, Sarah Brotman, and Kevin Currin for their guidance and help with eQTL analyses and Erika Deoudes for her graphic design contributions.

## Supplemental Information

**Supplemental Fig 1:**
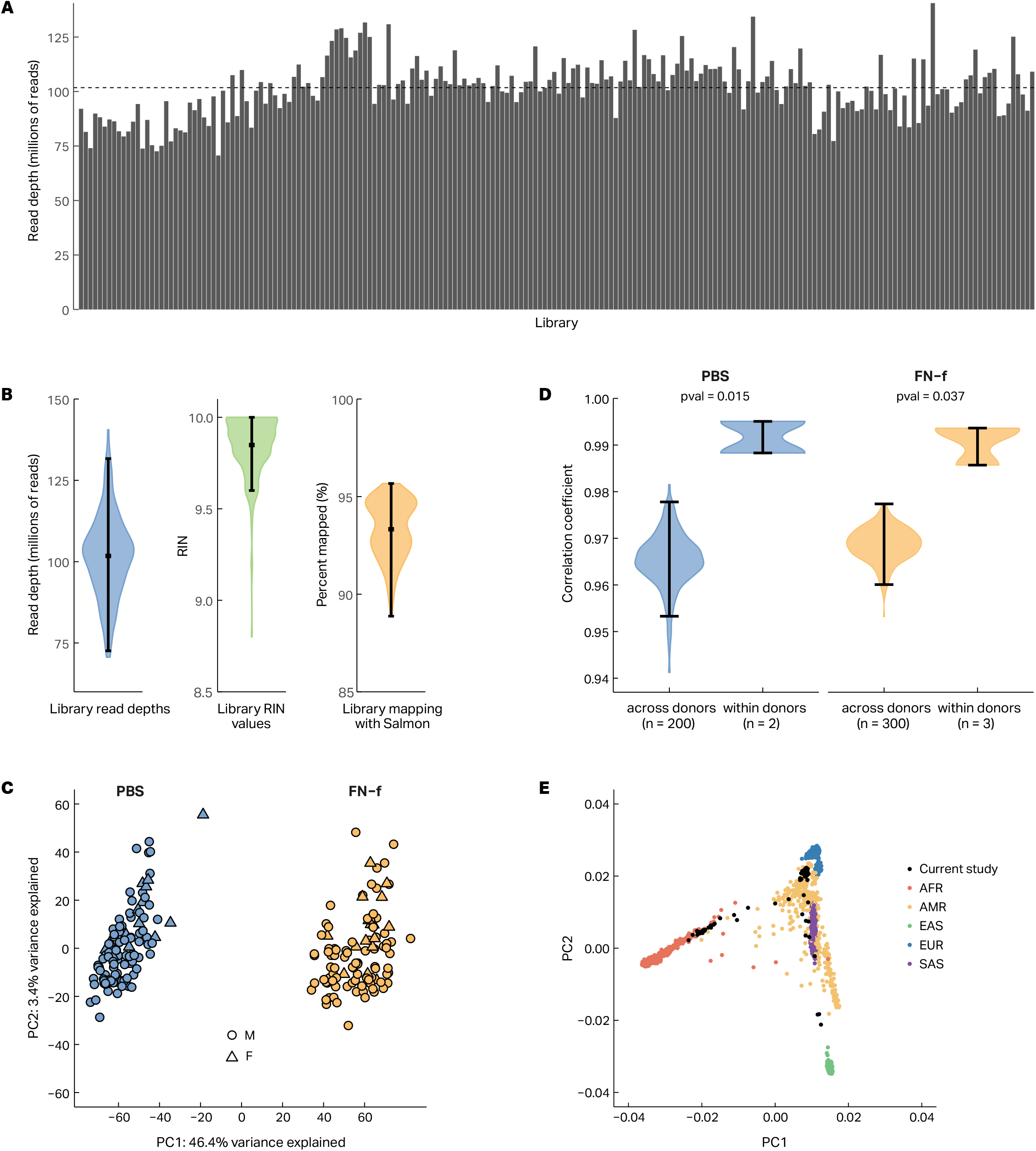
RNA-seq and eQTL QC analyses. (**A**) Barplot depicting read depths for all 202 RNA-seq data sets used in this study. The dashed line represents the mean library read depth (101.8 million reads). (**B**) Violin plots depicting the distributions of read depths (left), RIN scores (middle), and mappability (right) of RNA-seq data sets used in this study. (**C**) The correlation of gene expression between vs within donors for PBS and FN-f samples is visualized via violin plots. (**D**) PCA plots reveal that RNA-seq samples cluster largely by treatment. Samples are colored by treatment and shaped by donor sex. (**E**) Principal component analysis of donor genotyping data calculated with EIGENSTRAT. Study data, colored in black, is overlaid with 1000 Genomes data, which are colored by superpopulations as denoted by the 1000 Genomes Project. AFR: African, AMR: Admixed American, EAS: East Asian, EUR: European, SAS: South Asian.

**Supplemental Fig 2:**
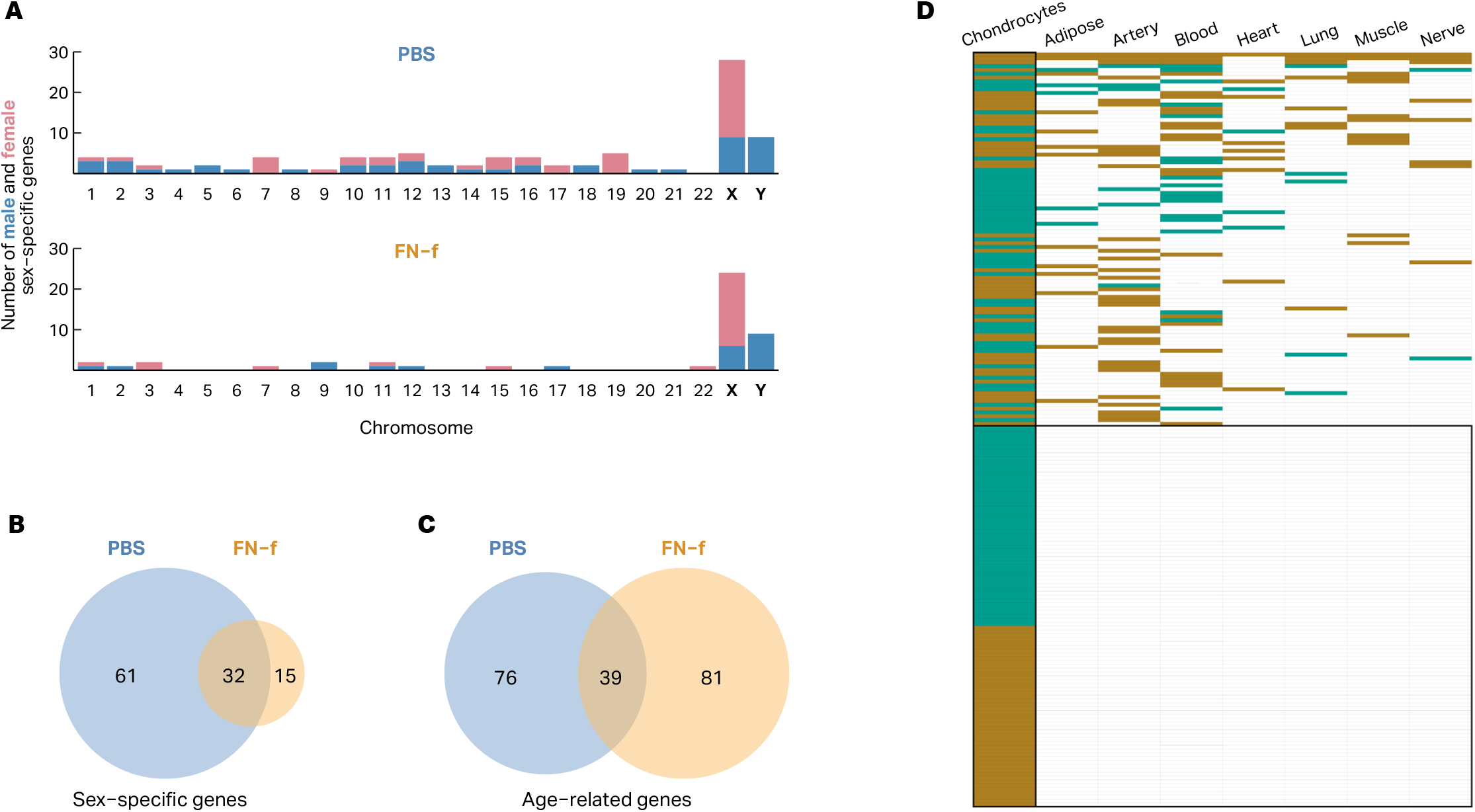
Overview of sex and age related gene expression differences. (**A**) A barplot showing that most, but not all, sex-biased genes in chondrocytes in both PBS and FN-f reside on X and Y chromosomes. Venn diagrams depicting the overlap of sex (**B**) and age (**C**) biased genes between conditions.

**Supplemental Fig 3:**
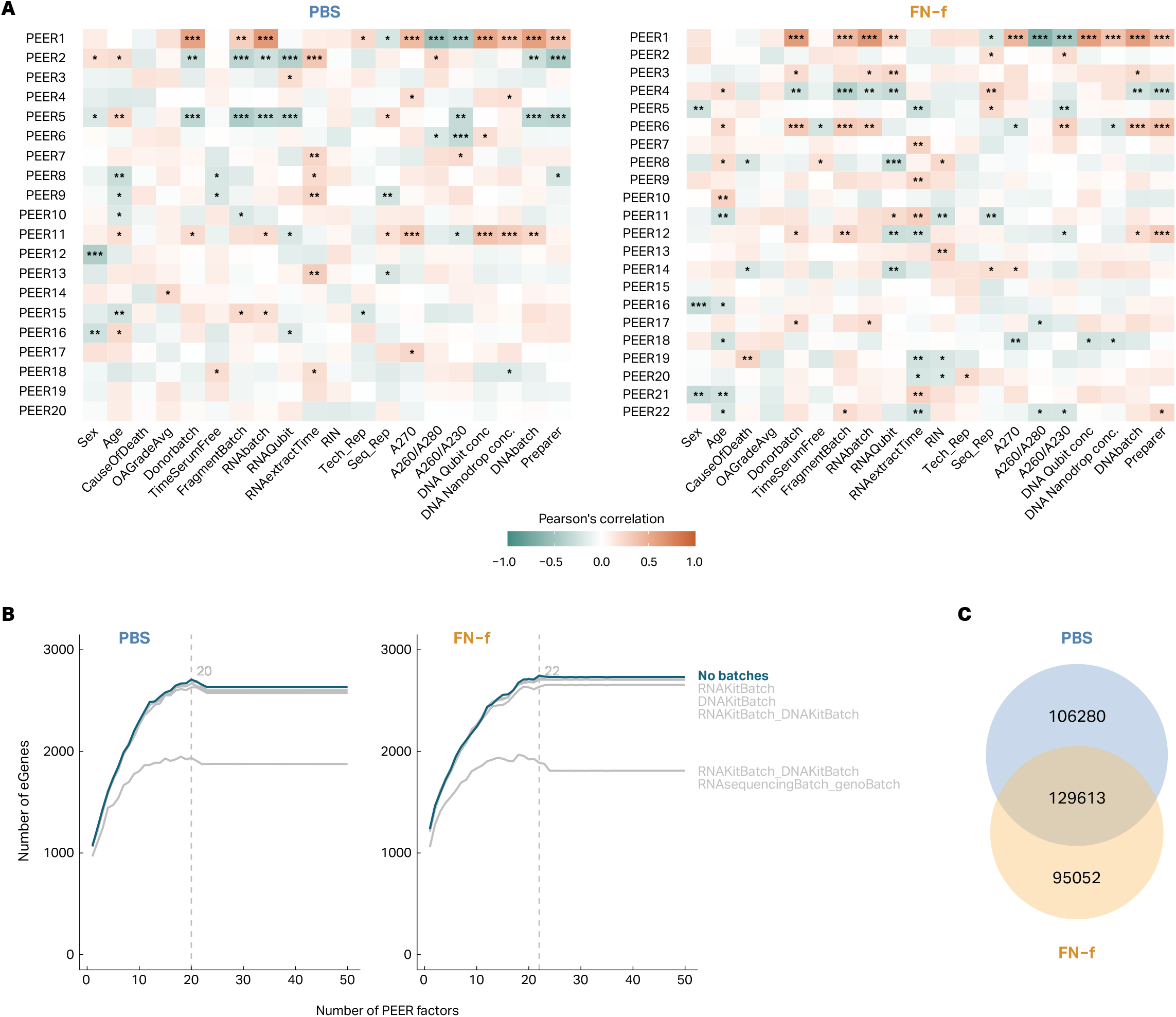
Optimization of eQTL discovery. (**A**) Heatmaps of Pearson’s correlations between calculated PEER factors and known technical covariates in PBS (left) and FN-f (right) samples. * p-value < 0.05, ** p-value < 0.01, *** p-value < 0.001. (**B**) Number of significant eGenes as a result of correcting for 1-50 PEER factors and including various additional batch covariates. 20 PEER factors with no additional batches and 22 PEER factors with no additional batches yielded the most significant eGenes in PBS (left) and FN-f (right) with eQTL mapping. (**C**) Venn diagram of all significant eQTL-eGene pairs identified in PBS and FN-f.

**Supplemental Fig 4:**
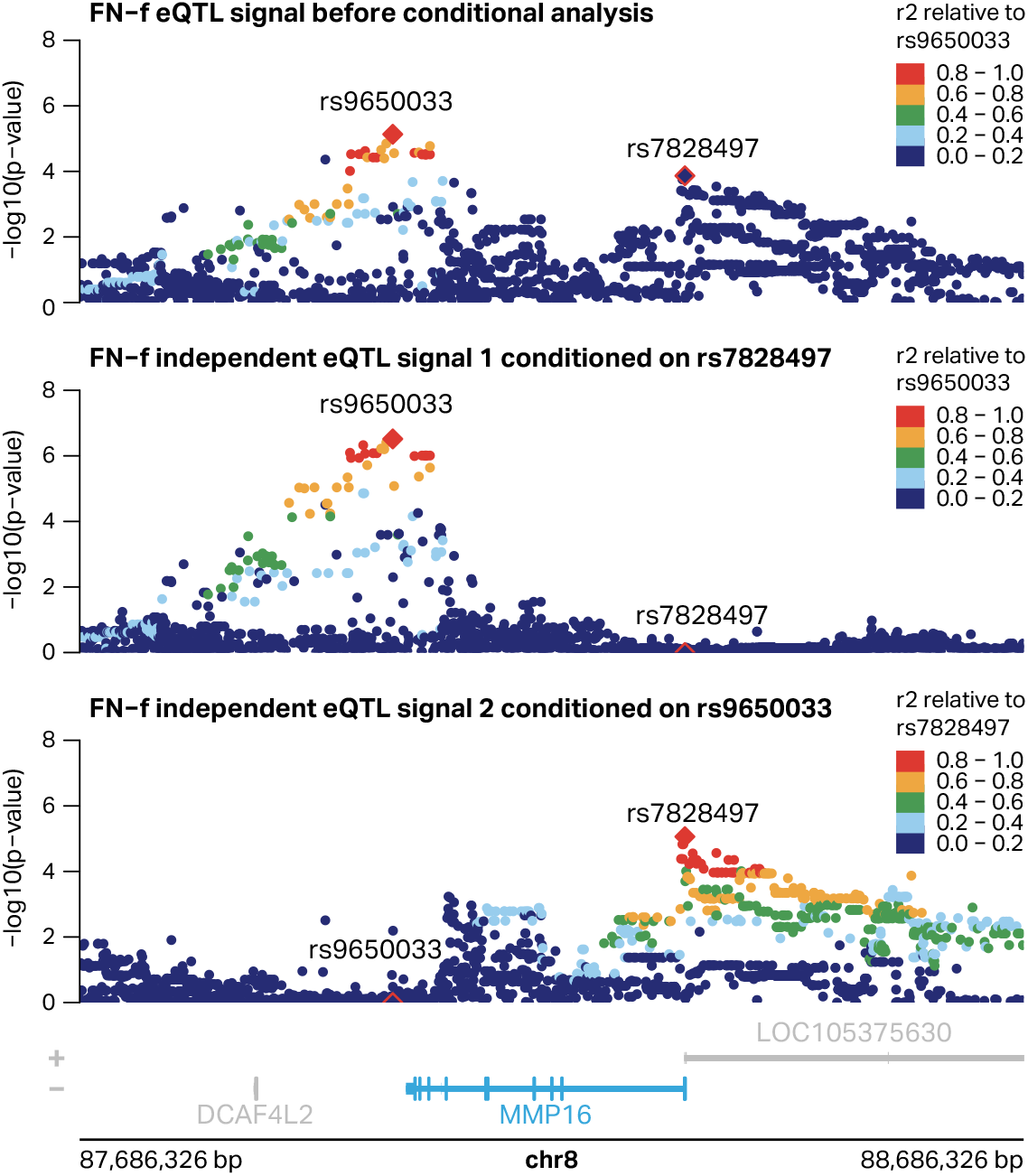
Conditional eQTL mapping identifies secondary signals. Locus zooms of the FN-f MMP16 eQTL signal before conditional analysis and after isolating independent signals. rs9650033 and rs7828497 are lead variants for the independent signals and are not in LD with each other. Each signal is colored by LD relative to the signal lead variant.

**Supplemental Fig 5:**
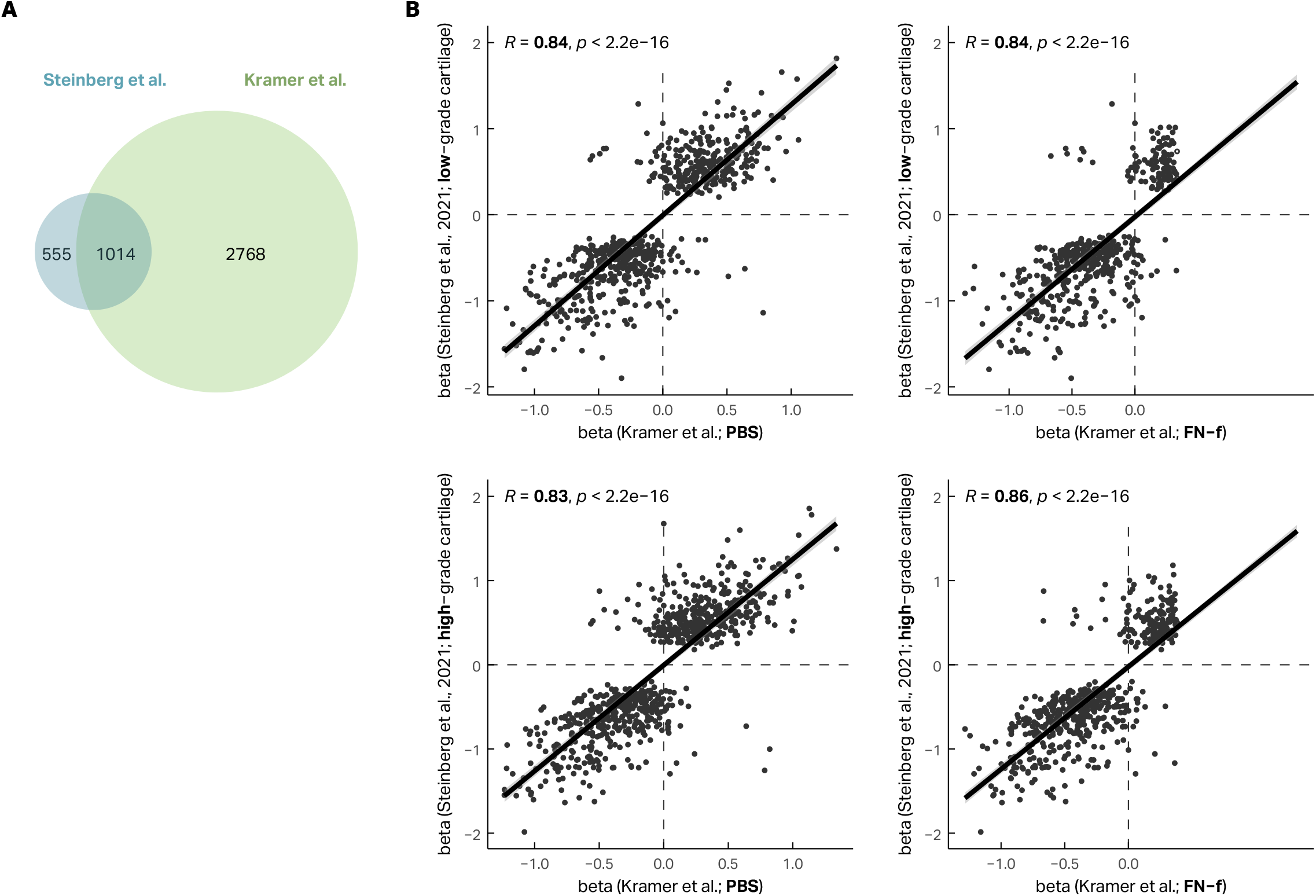
Comparison of chondrocyte eQTLs to those identified by Steinberg et al. (**A**) A Venn diagram shows the overlap of eGenes between our study (PBS and FN-f) and Steinberg et al. (low-grade and high-grade cartilage). (**B**) Scatterplots comparing the effect sizes (beta) of Steinberg et al. lead variants for shared eGenes between the current study and Steinberg et al. Effect sizes are shown comparing PBS to low-grade cartilage (top left), PBS to high-grade cartilage (bottom left), FN-f to low-grade cartilage (top right), and FN-f to high-grade cartilage (bottom right) eQTLs. R represents the Pearson correlation coefficient between beta values.

**Supplemental Fig 6:**
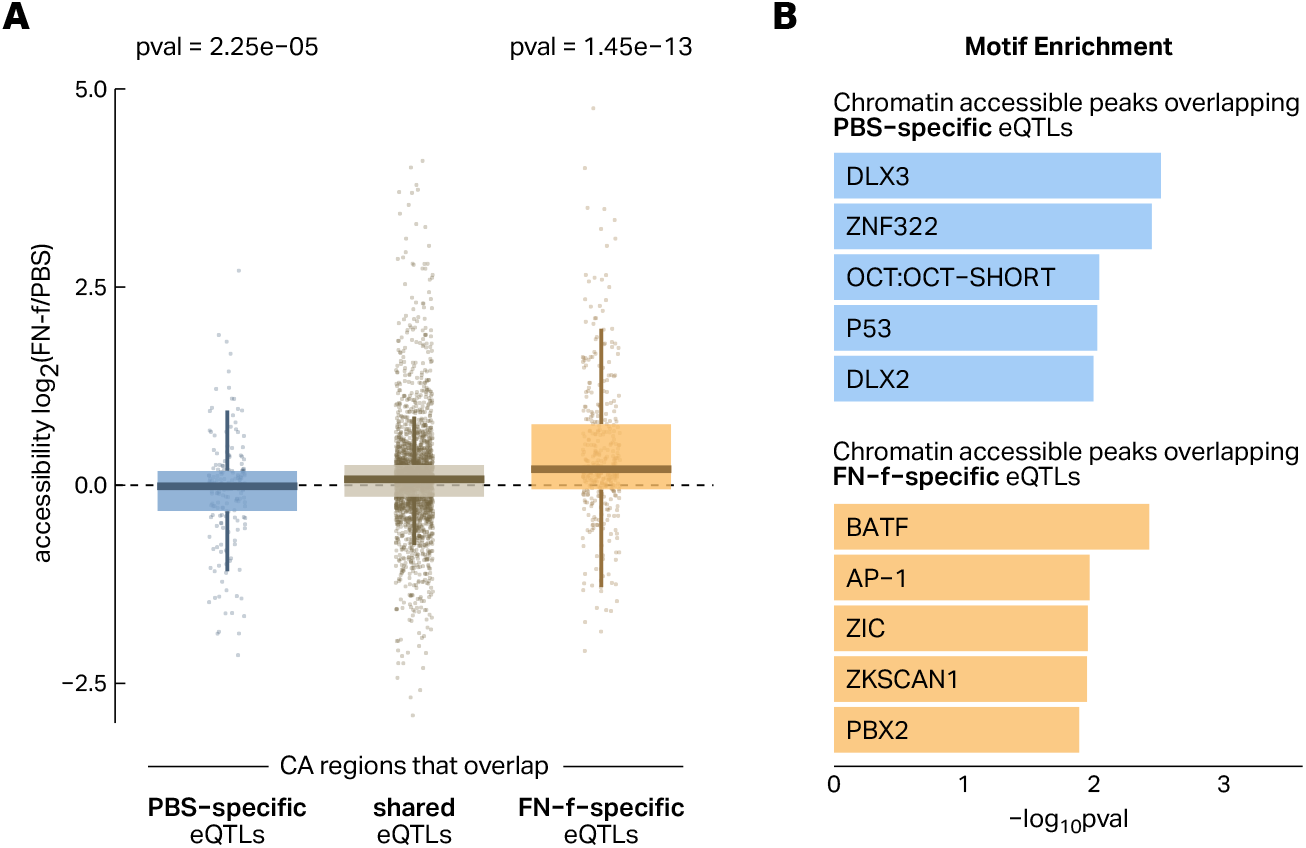
Chromatin accessibility supports condition-specific eQTLs. (**A**) Boxplots showing the log2 fold change in accessibility (FN-f/PBS) of chromatin-accessible (CA) regions overlapping high-confidence PBS-specific eQTLs (blue), shared eQTLs (tan), and high-confidence FN-f-specific eQTLs (yellow). Each set of eQTLs used to overlap with CAs contains the lead variant and any variants in high LD (> 0.8) with the lead. P-values are calculated from a Wilcox test. (**B**) Transcription factor motif enrichment for CAs overlapping PBS-specific (blue) or FN-f specific eQTLs (yellow).

**Supplemental Fig 7:**
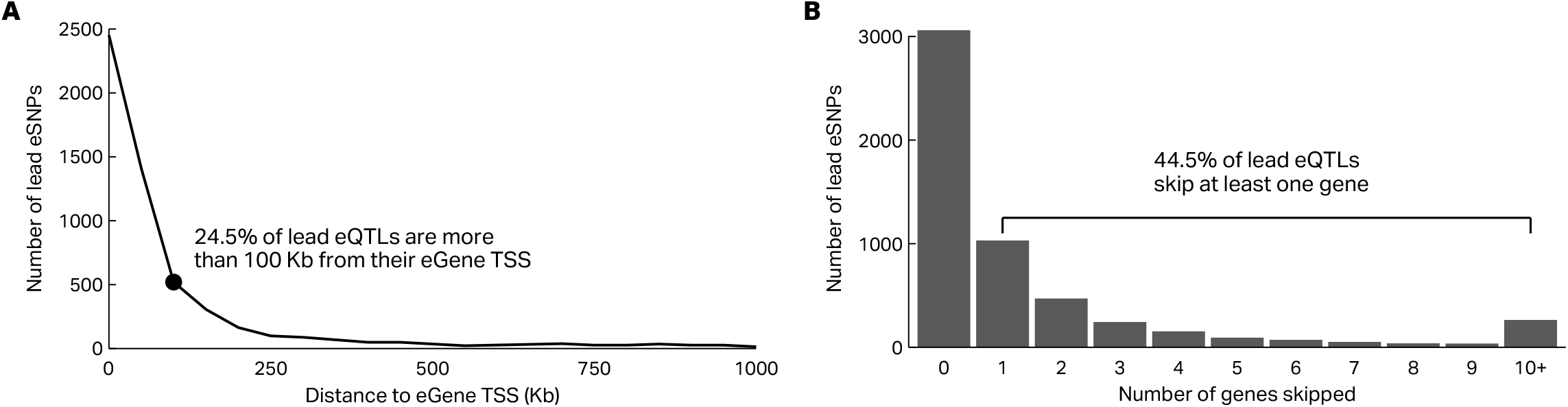
Many lead eQTLs suggest distal regulatory contacts with their eGenes. (**A**) Kilobase distance of PBS and FN-f lead eSNPs to their eGene transcription start sites (TSS) as defined in methods. (**B**) Barplot showing the number of other genes skipped between the assignment of a lead eSNP to its eGene.

**Supplemental Fig 8:**
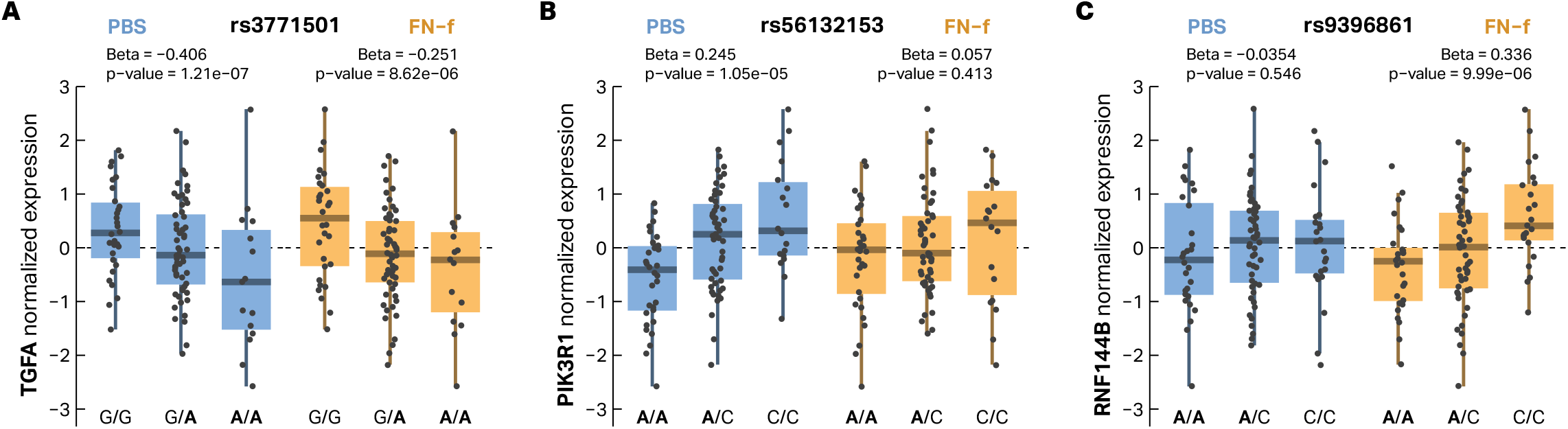
Effects of GWAS risk variants for examples of PBS-specific, FN-f-specific, and shared eQTL colocalizations. (**A**) The risk allele (A) of GWAS variant rs3771501 is associated with decreased expression of *TGFA* in both PBS and FN-f. (**B**) The risk allele (A) of GWAS variant rs56132153 is associated with decreased expression of *PIK3R1* only in PBS. (**C**) The risk allele (A) of GWAS variant rs9396861 is associated with decreased expression of *RNF144B* only after FN-f treatment. Boxplots depict donor genotypes at GWAS variants vs normalized gene expression with GWAS risk alleles bolded within labeled genotypes.

**Supplemental Fig 9:**
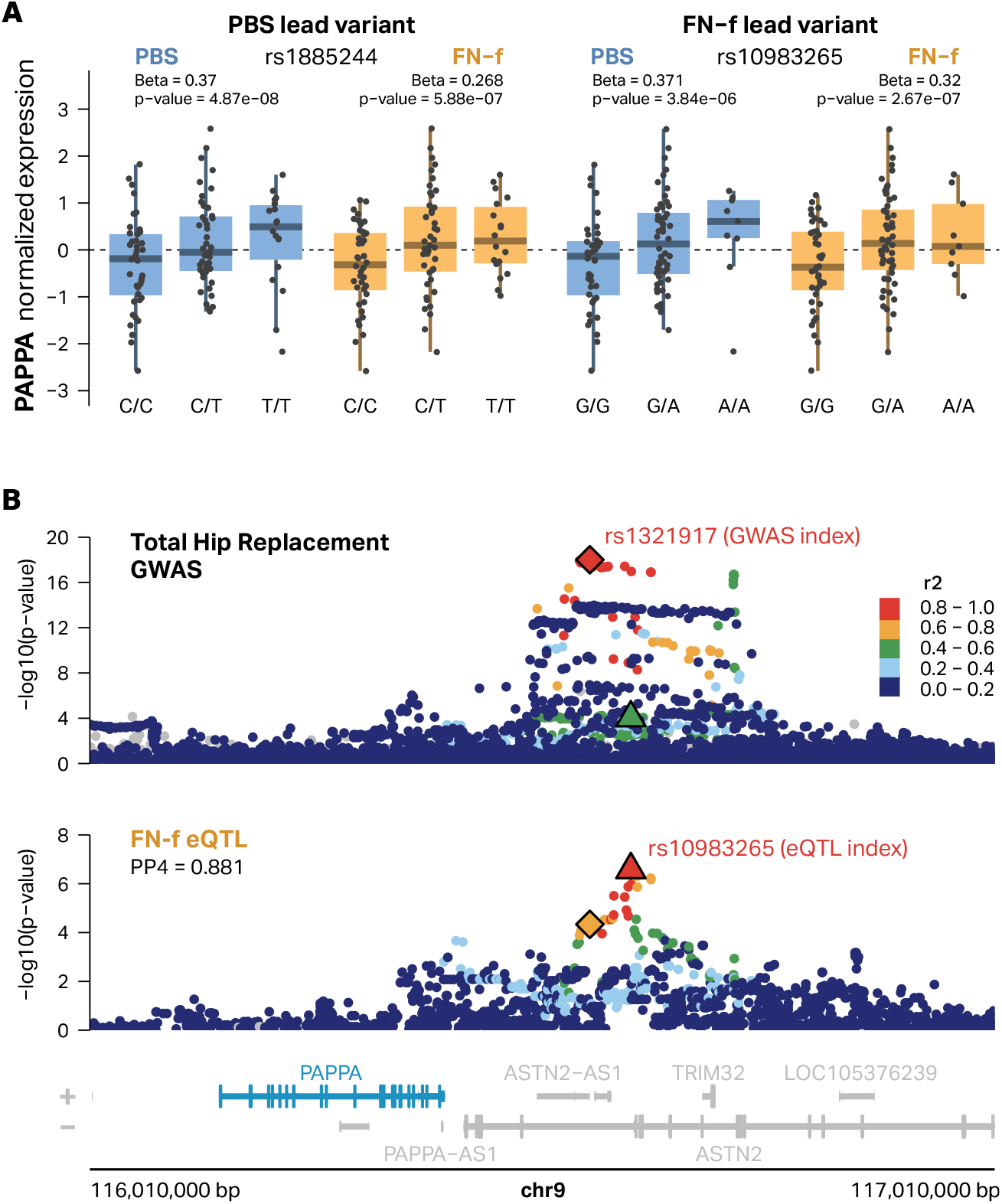
*PAPPA* eQTL variants in PBS and FN-f colocalize with OA GWAS. (**A**) Boxplots depicting the effects of the PBS lead eQTL rs1885244 (left) and FN-f lead eQTL rs10983265 (right) on *PAPPA* expression. Both lead variants show the same direction of effect in both PBS and FN-f. (**B**) Locus zoom depicting the colocalization between Total Hip Replacement GWAS and the eQTL signal identified in FN-f. Total Hip Replacement GWAS is colored by LD relative to the GWAS index rs1321917 according to the 1000 Genomes European reference panel.

**Supplemental Fig 10:**
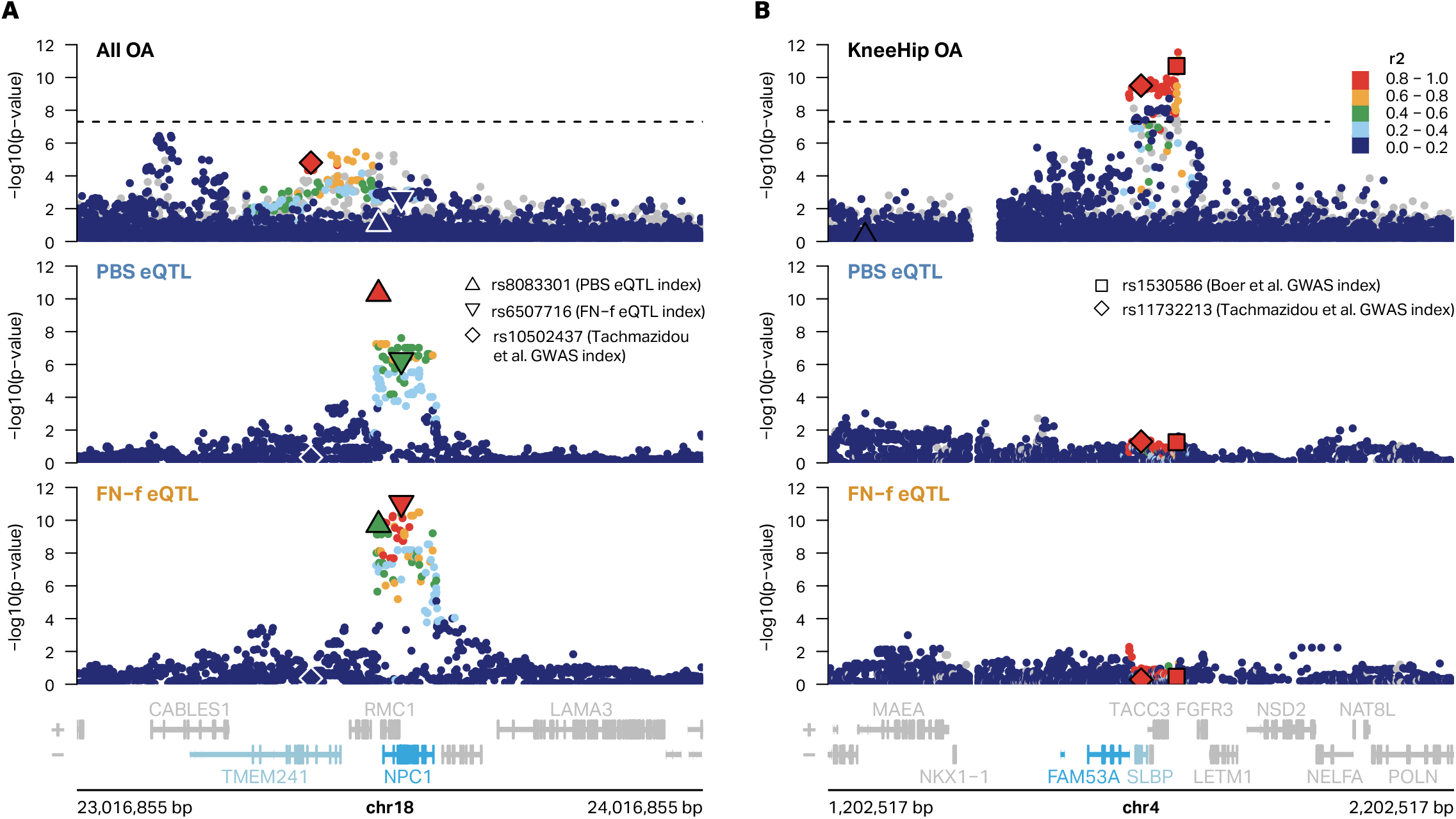
eQTL signals for *NPC1* and *FAM53A* do not colocalize with OA GWAS. (**A**) An All OA GWAS signal that was previously identified as colocalized with *NPC1* eQTL is identified as an eQTL in our study but is no longer significant in updated OA GWAS from Boer et al. Association plots depict Boer et al. All OA GWAS (top) and PBS (middle) and FN-f (bottom) eQTL signals for *NPC1*. All OA GWAS is colored by LD relative to the GWAS index rs10502437 from Tachmazidou et al. according to the 1000 Genomes European reference panel. PBS eQTL is colored by LD relative to PBS lead variant rs8083301 and FN-f eQTL is colored by LD relative to FN-f lead variant rs6507716. The Steinberg et al. lead variant for NPC1 (highlighted blue) resided in an intron of *TMEM241* (highlighted light blue). (**B**) A previously identified eQTL signal for *FAM53A* that colocalized with KneeHip OA GWAS is not identified in the current study. A KneeHip OA GWAS signal identified by Tachmazidou et al. remains significant in Boer et al. with the lead variants from both studies in high LD (> 0.8) according to the 1000 Genomes European reference panel (top). *FAM53A* PBS eQTL (middle) and FN-f eQTL (bottom) signals are not significant. Association plots are all colored by LD relative to the Boer et al. GWAS index variant rs1530586 according to the 1000 Genomes European reference panel. The Steinberg et al. lead variant for *FAM53A* (highlighted blue) resided in an intron of *SLBP* (highlighted light blue).

