## Supplemental Tables S1-14 Captions for "Response eQTLs, chromatin accessibility, and 3D chromatin structure in chondrocytes provide mechanistic insight into osteoarthritis risk"

### Supplemental Table Captions:

**Supplemental Table 1.** Genes differentially expressed in response to FN-f treatment.
The direction of response is indicated with "-", meaning downregulated, and "+", meaning upregulated. The most significant genes with the largest effects (adjusted p-value < 0.01 and absolute log2 fold change > 2) are denoted with "---" and "+++".
symbol: gene symbol
gene_id: gene ENSEMBL ID
padj: adjusted p-value
log2FoldChange: log2 fold change of FN-f expression relative to PBS
FNF response: direction of gene response to FN-f

**Supplemental Table 2.** All GO terms and KEGG pathways that are enriched in FN-f upregulated and downregulated genes.
Tables are split by GO/KEGG and Upregulated/Downregulated.
GO term columns:
TermID: GO term ID
Term: GO term name
parentTermID: GO parent term ID as determined by string similarity
parentTerm: GO parent term name as determined by string similarity
Enrichment: enrichment score
-log10pval: -log10 p-value of the GO term enrichment
Genes in Term: number of genes in that GO term
Target Genes in Term: number of genes in that term from the gene set of interest
Fraction of Targets in Term: Target Genes in Term/Total Target Genes
Total Target Genes: Number of target genes tested. This number will be the same within each dataset.
Total Genes: Number of total genes used for GO enrichment analysis. This number will be the same within each dataset.
Gene Symbols: The gene symbols of all the target genes identified with a GO term.

KEGG pathway columns:
TermID: KEGG pathway ID
Term: KEGG pathway name
Enrichment: enrichment score
-log10pval: -log10 p-value of the KEGG pathway enrichment
Genes in Term: number of genes in that KEGG pathway
Target Genes in Term: number of genes in that term from the gene set of interest
Fraction of Targets in Term: Target Genes in Term/Total Target Genes
Total Target Genes: Number of target genes tested. This number will be the same within each dataset.
Total Genes: Number of total genes used for KEGG enrichment analysis. This number will be the same within each dataset.
Gene Symbols: The gene symbols of all the target genes identified with a KEGG pathway.

**Supplemental Table 3.** Sex-specific genes.
symbol: gene symbol
gene_id: ENSEMBL gene ID
sex_padj_PBS: adjusted p-value of sex-specific gene in PBS
sex_log2FoldChange_PBS: log2 fold change (M/F) of gene in PBS
sex_padj_FNF: adjusted p-value of sex-specific gene in FN-f
sex_log2FoldChange_FNF: log2 fold change(M/F) of gene in FN-f
M/F: column indicating whether the gene is male-biased (M) or female-biased (F)
FNF response: direction of response of sex-specific gene to FN-f treatment. A missing value indicates the gene was not differentially expressed with FN-f. "+++": high-effect FN-f upregulation; "+": FN-f upregulation; "---": high-effect FN-f downregulation; "-": FN-f downregulation.
Change with OA: direction of expression of sex-specific gene in OA tissue. A missing value indicates the gene was not significantly expressed in OA.
OA study: the OA study the sex-specific gene was identified in, if applicable.

**Supplemental Table 4.** Sex-specific gene expression in multiple tissues.
This table corresponds to the heatmap in Fig 2B.
tissue_no: The number column a tissue corresponds to in the heatmap, from left to right
tissue: tissue/cell type name
gene_no: The number row a gene corresponds to in the heatmap, from top to bottom
sex: The sex the gene is associated with.
effsize: Effect size, log2 fold change
effsize_se: Standard error of effect size

**Supplemental Table 5.** Age-related genes.
symbol: gene symbol
gene_id: gene ENSEMBL ID
age_padj_PBS: adjusted p-value of age LRT in PBS
age_padj_FNF: adjusted p-value of age LRT in FN-f
Change with age: direction of expression change with older donors
FNF response: direction of response of age-related gene to FN-f treatment. A missing value indicates the gene was not differentially expressed with FN-f. "+++": high-effect FN-f upregulation; "+": FN-f upregulation; "---": high-effect FN-f downregulation; "-": FN-f downregulation.
Change with OA: direction of expression of age-related gene in OA tissue. A missing value indicates the gene was not significantly expressed in OA.
OA study: the OA study the age-related gene was identified in, if applicable.

**Supplemental Table 6.** All GO terms that are enriched in genes that increase and decrease in expression with age.
Tables are split by Up with Age/Down with Age.
TermID: GO term ID
Term: GO term name
parentTermID: GO parent term ID as determined by string similarity

parentTerm: GO parent term name as determined by string similarity
Enrichment: enrichment score
-log10pval: -log10 p-value of the GO term enrichment
Genes in Term: number of genes in that GO term
Target Genes in Term: number of genes in that term from the gene set of interest
Fraction of Targets in Term: Target Genes in Term/Total Target Genes
Total Target Genes: Number of target genes tested. This number will be the same within each dataset.
Total Genes: Number of total genes used for GO enrichment analysis. This number will be the same within each dataset.
Gene Symbols: The gene symbols of all the target genes identified with a GO term.

**Supplemental Table 7.** PBS and FN-f lead eQTLs and eGenes.
Conditionally independent lead eSNP-eGene pairs, separated by condition.
gene_id: eGene ENSEMBL ID
gene_name: eGene name
gene_chr: gene chromosome
gene_tss: gene transcription start site used for eQTL mapping, as defined in Methods
gene_end: gene end
gene_strand: gene strand
num_cis_variants: number of genetic variants that were *in cis* and tested for a gene
rsID: lead eSNP rsID
variantID: lead eSNP variant ID in the format “variant chromosome:variant position: allele 1: allele 2”
variant pos: lead eSNP position
beta: beta estimate of effect size
beta_se: standard error of calculated beta
nom_pval: nominal p-value
minor_allele: the minor allele, also the assessed or effect allele, for the lead eSNP. Beta estimates are relative to this allele.
MAF: minor allele frequency
eGene_nominal_threshold: the local nominal p-value significance threshold for an eGene
eGene_qval: the globally adjusted q-value for an eGene based on permutation testing
signal: iteration of conditional analysis. 0 = primary signal, 1 = secondary signal, etc.

**Supplemental Table 8.** Study comparisons to Steinberg et al.
eGenes and lead variants:
Comparison of eGenes and lead variants between Steinberg et al. low grade and high grade cartilage eQTL datasets and current study PBS and FN-f eQTL datasets. Steinberg et al. variant coordinates were lifted over to hg38 to match the current study. All variant IDs are in the format “chr:hg38pos:ref:alt”.

gene_id: eGene ENSEMBL ID
gene_symbol: eGene name

variantID_Kramer_PBS: lead variant ID for primary eGene signal in PBS eQTL dataset
variantID_Kramer_FNF: lead variant ID for primary eGene signal in FN-f eQTL dataset
variantID_Steinberg_lowgrade: lead variant ID for eGene in Steinberg low grade cartilage dataset
variantID_Steinberg_highgrade: lead variant ID for eGene in Steinberg high grade cartilage dataset
study: Which study the eGene was identified in: Kramer, Steinberg, or both

Study Designs:
Overview of current eQTL study and Steinberg et al. OA eQTL study datasets, including sample demographics, cartilage joint site, RNA-sequencing library preparation, genotyping platform, quality control, SNP imputation, eQTL calling methods, and colocalization methods.
HWE, Hardy-Weinberg Equilibrium; MAF, minor allele frequency; HRC, Haplotype Reference Consortium; PC, principal component; FDR, false discovery rate; GWAS, genome-wide association study; PP4, posterior probability 4 obtained from coloc

**Supplemental Table 9.** Condition-specific eQTLs in PBS and FN-f.
All lead eSNP-eGene pairs with a significant genotype by condition interaction effect. High confidence PBS-specific and FN-f-specific eQTLs are denoted by the "high_conf" column.

gene_id: eGene ENSEMBL ID
gene_name: eGene name
gene_chr: gene chromosome
gene_tss: gene transcription start site used for eQTL mapping, as defined in Methods
gene_end: gene end
gene_strand: gene strand
rsID: lead eSNP rsID
variantID: lead eSNP variant ID in the format "variant chromosome:variant position: allele 1: allele 2"
variant pos: lead eSNP position
PBS_beta: beta estimated effect size of lead eSNP-eGene in PBS
PBS_beta_se: standard error of calculated beta in PBS
FNF_beta: beta estimated effect size of lead eSNP-eGene in FN-f
FNF_beta_se: standard error of calculated beta in FN-f
minor_allele: the minor allele, also the assessed or effect allele, for the lead eSNP. Beta estimates are relative to this allele.
MAF: minor allele frequency
interaction_pval: p-value from ANOVA testing of genotype by condition interaction
high_conf: a column indicating whether the eQTL was high confidence condition-specific. A missing value indicates it is not part of this group.

**Supplemental Table 10.** KEGG pathways enriched in high-confidence PBS-specific and high-confidence FN-f-response eGenes.

TermID: KEGG pathway ID

Term: KEGG pathway name
Enrichment: enrichment score
-log10pval: -log10 p-value of the KEGG pathway enrichment
Genes in Term: number of genes in that KEGG pathway
Target Genes in Term: number of genes in that term from the gene set of interest
Fraction of Targets in Term: Target Genes in Term/Total Target Genes
Total Target Genes: Number of target genes tested. This number will be the same within each dataset.
Total Genes: Number of total genes used for KEGG enrichment analysis. This number will be the same within each dataset.
Gene Symbols: The gene symbols of all the target genes identified with a KEGG pathway.

**Supplemental Table 11.** Differential accessible peaks.

chrom: chromosome
start: peak start
end: peak end
width: peak width
baseMean: base mean of peak counts
log2FoldChange: log2 fold change of accessibility in FN-f relative to PBS
lfcSE: standard error of log2 fold change
padj: adjusted p-value

**Supplemental Table 12.** Differential loops.

chrom1: chromosome of loop anchor 1
start1: start of loop anchor 1
end1: end of loop anchor 1 (10 kb from start1)
chrom2: chromosome of loop anchor 2
start2: start of loop anchor 2
end2: end of loop anchor 2 (10 kb from start2)
log2FoldChange: log2 fold change of loop contact frequency in FN-f relative to PBS
lfcSE: standard error of log2 fold change
padj: adjusted p-value
cluster: looping cluster, either gained or lost with FN-f treatment
DonorX_XXX_rX_raw: raw loop counts for a given donor sample (1-4), replicate (r1 or r2), and treatment condition (PBS or FNF)

**Supplemental Table 13**. Extended characterization of colocalizations between PBS and FN-f eQTL and OA GWAS signals.
OA, osteoarthritis; GWAS, Genome-wide association study; THR, Total Hip Replacement; OR, odds ratio; PBS coloc PP4, posterior probability 4 obtained from coloc testing with the PBS signal for an eQTL; FN-f coloc PP4, posterior probability 4 obtained from coloc testing with the FN-f signal for an eQTL; M vs F, male vs female

**Supplemental Table 14.** Donor characteristics for all collected datasets.
Ancestries for eQTL-related donors (i.e. genotyping and RNA-seq) were determined from principal component analysis with 1000 Genomes samples using EIGENSTRAT. Definitions of ancestry superpopulations are defined by 1000 Genomes.
AFR, African; AMR, Admixed American; EAS, East Asian; EUR, European; SAS, South Asian
